# A metagenomic thermostable monomeric meganuclease with novel specificity and unique palindromic 3’ overhangs

**DOI:** 10.64898/2026.03.18.712669

**Authors:** Gol Mohammad Dorrazehi, Matthew Penner, Christina Athanasiou, Lise Boursinhac, Juan Carlos Mobarec, Carl Webster, Monika Papworth, Florian Hollfelder

## Abstract

As an alternative to historical enzyme isolation, metagenomic databases (e.g. MGnify) provide information on vast unculturable microbial diversity, especially from extreme environments, and constitute an enormous source of functional proteins. Conservative mining of these data by close sequence homology alone tends to identify merely different versions of known enzymes. Here we present a discovery strategy of meganucleases based on wider capture of less homologous enzymes with new function in metagenomic databases, incorporating metadata with homology, relying on cell-free expression to bypass host incompatibility and the need for purification, along with using deep sequencing for experimental assessment of substrate specificity and cleavage pattern, circumventing classical gel-based profiling. Specifically, we discovered the temperature-stable (>55°C), intron-encoded LAGLIDADG meganuclease I-MG11 that recognizes a 17 base pair sequence to generate unique 4 base pair palindromic 3′-overhangs — the first monomeric meganuclease to produce such overhangs. Co-folding models of I-MG11 bound to DNA provide a structural context for enzyme-DNA interactions, highlighting differences from other monomeric LAGLIDADG meganucleases (e.g. I-SceI) shaped by InDels (insertion-deletions) in the DNA binding region that may cause specificity changes. Our strategy streamlines *bona fide* identification and annotation of meganucleases, while the unique properties of I-MG11 expand the molecular biology toolbox.

**GRAPHICAL ABSTRACT:** 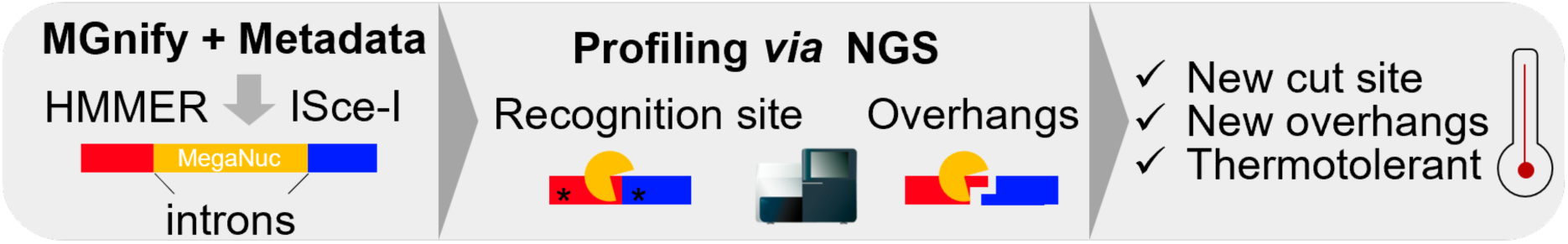

## INTRODUCTION

Meganucleases (also called homing endonucleases) offer unique advantages for precise genomic and *in vivo* DNA manipulation (1–4), notwithstanding the advent of newer easily programmable genome editing tools, mainly derived from clustered regularly interspaced short palindromic repeats-Cas (CRISPR-Cas) technology (5–7). Their inherent high specificity of meganucleases and their ability to induce highly efficient homologous recombination in a one-component reaction makes them useful, complementary reagents in any gene editing application, without the need to add a separate guide RNA (as in CRISPR-Cas) or requiring two proteins that must dimerize (as in zinc finger nucleases (ZFNs) and transcription activator-like effector nucleases (TALENs)). Such compact, all-in-one architecture can simplify meganuclease delivery (*via* plasmids or mRNA) into target cells, especially for viral vectors where cargo capacity is a concern (8). Despite these inherent advantages, the long target DNA sequences (14-40 base pairs) recognised by meganucleases are harder to find, as they are statistically much less likely to occur randomly in a large genome like a human’s (9–12). However, reprogramming of a meganuclease for a new target or specific applications (13–15) is more challenging, because the convenient modular manipulation via the external guide sequence is unavailable. Instead, the entire enzyme has to be engineered or new enzymes have to be found to make a larger thesaurus of enzymes with specific cleavage patterns available (8,16). Here, we set out to discover novel meganucleases as a basis for further enzyme engineering and functional improvements.

The traditional methods of enzyme discovery, primarily relying on detailed elucidation of biological mechanisms in culturable microorganisms, have significant limitations. It is estimated that only a small fraction (less than 1%) of microbial diversity can be cultivated in laboratory settings (16–18) leaving an immense, untapped reservoir of enzymatic potential hidden within uncultured microbes. The advent of metagenomics has revolutionized the field of enzyme discovery, because the direct extraction and analysis of genetic material from environmental samples to bypass the need for microbial cultivation. The availability of vast metagenomic data offers an unprecedented opportunity to access this vast natural genetic diversity (19, 20). Access to collective genomes of entire microbial communities provides a route to a wealth of genes encoding novel enzymes but the challenge of assignment of function has to be met to make use of them based on bona fide annotation.

Among the various metagenomic resources available, databases like MGnify (https://www.ebi.ac.uk/metagenomics) (21, 22) stand out as critical platforms for accelerating enzyme discovery, especially for identification of closely related enzymes. MGnify provides comprehensive, assembled and annotated metagenomic and metatranscriptomic data from diverse environments and constitutes the largest public database of open reading frames available and its > 2 billion entries to-date offer an untapped reservoir of functional proteins (22, 23). By leveraging such unprecedented information, scientists can employ homology searches or more complex algorithmic exploration (24) to identify candidates for experimental screening. Complementary to these approaches, functional metagenomic screens can overcome intrinsically conservative homology-guided searches, revealing functional annotations in unexplored (25, 26) or shallow (24, 27) sequence contexts that can subsequently be improved by directed evolution (26, 28). The reciprocal interplay of bioinformatics and experimental screening will accelerate bioprospecting of useful enzymes for applied biocatalysis and biotechnology.

In this work we designed a fast workflow to discover and characterise novel meganucleases. We first harvested data from the MGnify database and then characterised the recognition sequence, cleavage pattern, specificity and thermostability of a candidate enzyme using cell free protein expression and deep sequencing. This method allowed us to fully characterise a newly discovered meganuclease in only a few days, and could easily be scaled up to multiple enzymes. We successfully applied it to discover the monomeric (single-chain) meganuclease I-MG11 from the LAGLIDADG family. I-MG11 is a thermostable enzyme from an extreme marine hydrothermal environment that recognizes a 17 base pair target sequence for cleavage and generates four base pair palindromic 3’-overhangs.

## MATERIAL AND METHODS

### Extraction of LAGLIDADG meganuclease enzymes from MGnify

The protein sequence of the I-SceI meganuclease was used as query bait to retrieve sequences from MGnify (https://www.ebi.ac.uk/metagenomics) (21, 22), going further than a previous, relatively conservative homology search (28% vs 57%) (29). Based on ranking by E-value, the top 50 hits were used to generate a Maximum Likelihood (ML) phylogenetic tree (**Figure S1**; sequences available in **Supplementary File S1**). Subsequently, 16 random sequences were selected from across this tree for further cloning.

### Cloning and expression

Based on protein sequences retrieved from MGnify, *E. coli* codon-optimized genes were produced by GeneArt Gene Synthesis (Thermo Fisher Scientific). The genes were cloned into pET24a plasmid using NEBuilder assembly (New England Biolabs), using the complementary overhangs included in the genes during synthesis, and introduced in the plasmid backbone by primers during the amplification. Assembly mix was transformed into either *E. coli* HST08 Stellar (Takara Bio) or *E. coli* NEBExpress^®^ Iq (NEB) competent cells.

### Protein expression and purification

*In vitro expression:* Genes cloned under control of a T7 promoter (in a pET24a plasmid) were expressed using the PURExpress^®^ In Vitro Protein Synthesis Kit (NEB) according to manufacturer’s instruction.

*E. coli expression and purification of I-MG11:* For tighter regulation the pBAD Expression System (Thermo Fisher Scientific) was used, which contains an L-arabinose inducible promoter (P_BAD_). A strep tag II sequence (WSHPQFEK) proceeding a GlyAlaGly (GAG) linker was added to the C-terminus of I-MG11 by amplifying the *E. coli* optimized gene from the pET24a plasmid using the following primers (with overlapping regions shown in lower case):

*Fw_MG11-gene:*

5’_ccgtttttttgggctaacaggaggaattaaccATGTTTCTGCGTGGTAAAAAGC_3’

*Rv_MG11-gene:*

5’_ attatttttcgaactgcgggtggctccaacctgCACCGTTACGCGGAAACGGCAGTTTG_3’

For pBAD plasmid backbone amplification following primers were used:

*Fw_CterStrep_pBAD:*

5’_aggttggagccacccgcagttcgaaaaataatGAGCTTGGCTGTTTTGGCGGATG_3’

*Rv_pBAD:*

5’_ggttaattcctcctgttagcc_3’

The PCR products were cloned into a pBAD plasmid backbone using NEBuilder assembly (New England Biolabs) and transformed into *E. coli* TOP10 competent cells, followed by verification by colony PCR and Sanger sequencing. To facilitate protein expression, from a single colony a starter culture was inoculated in media containing 5% Glucose and appropriate antibiotic, and incubated overnight at 37°C. Next day, a 1 litre culture was inoculated and protein expression was induced by addition of L-arabinose (0.5% v/v), followed by incubation at 20°C (16 hours). Cells were collected by centrifugation and lysed using sonication, and from soluble fraction proteins were purified using Strep-Tactin^®^ 4Flow^®^ high capacity FPLC column (IBA Lifesciences). Binding was carried out in wash buffer (100mM Tris/HCl, pH 8, 400 mM NaCl) and elution in wash buffer supplemented with 50 mM D(+)-biotin.

### Cloning of I-MG11 into plasmid

The putative 30 bp and central 20 bp targets were cloned into a pUC plasmid containing a kanamycin resistant gene to give constructs pUC-MG11-30bp and pUC-MG11-20bp, respectively. This was implemented following the QuickLib method (30) using the following primers (with overlapping regions shown in lower case and underlined):

*MG11_30_bp_target_in_pUC_fw:*

5’_*gttcttcgcccaccccagcttcaaaaTAAAAGGGTACGCGAGCTGGGTTCAAACCGgcgctctgaagttcctag_*3’

*MG11_20_bp_target_in_pUC_fw:*

*5’_***gttcttcgcccaccccagcttcaaaaGGGTACGCGAGCTGGGTTCAgcgctctgaagttccta g_***3’*

*MG11_target_in_pUC_rv:* 5’_**ttttgaagctggggtggg_**3’

### Amplification of linear Cy5-linked target

From the pUC-MG11-30bp plasmid, linear and fluorophore-linked targets were PCR-amplified using one of the primers labelled on the 5’ end with a Cy5 (Cyanine5) fluorophore:

*Cy5_Fw_target: Cy5_5’_TGGAGAATGGGAGGTTTTCTGGG_3’*

*Rv_target: 5’_TTCGTGCACACAGCCCAGCTTGG_3’*

### Nuclease Cleavage Site and Overhang Identification

#### NGS-based Nuclease Cleavage Site and Overhang Identification (NuCSOI)

The two circular plasmids (pUC-MG11-30bp and pUC-MG11-20bp) were used to assess nuclease cleavage sites with varying recognition region lengths: a 30 bp recognition sequence (5′-TAAAAGGGTTACGCGAGCTGGGTCAAACCG-3′) and a central 20 bp recognition sequence (5′-GGGTTACGCGAGCTGGGTCA-3′; derived by deleting 5 bp from each end of the 30 bp region).

After cleavage of plasmids by IVTT-expressed I-MG11, followed by enzymatic removal of overhangs using NEBNext^®^ Ultra™ II End Repair/dA-Tailing Module (New England Biolabs), samples were subjected to paired-end sequencing for single-base resolution cleavage site mapping. Sequencing generated 182,552 read pairs (30 bp construct) and 230,164 read pairs (20 bp construct). Reads were quality filtered with fastp v0.23.2 (Q30 threshold; automatic adapter trimming) and aligned to their respective circular plasmid references using BWA-MEM v0.7.17 (-M flag) with custom circular reference handling. Post-alignment filtering retained reads with MAPQ ≥30 and aligned length ≥50 bp; duplicates were not removed.

After filtering, the 30 bp dataset yielded 735 high-quality alignments (mean length: 154 bp; MAPQ: 60.0) and the 20 bp dataset yielded 595 alignments (mean length: 156 bp; MAPQ: 60.0). Coverage was computed at single-nucleotide resolution using the NuCSOI v1.0.0 Python pipeline, and cleavage sites were inferred from abrupt local coverage changes indicative of nuclease activity.

#### Run-off sequencing for cleavage site and overhang identification

Plasmid pUC-MG11-30bp was incubated with IVTT-expressed I-MG11 and the linearized plasmid was isolated by gel purification using QIAquick Gel Extraction Kit (QIAGEN), to minimize contamination of residual non-cleaved plasmids. Run-off Sanger sequencing using following primers that that anneal upstream and downstream to the I-MG11 target site in the pUC-MG11-30bp plasmid:

*Fw_target: Cy5_5’_TGGAGAATGGGAGGTTTTCTGGG_3’*
*Rv_target: 5’_TTCGTGCACACAGCCCAGCTTGG_3’*

### Nuclease substrate profiling by deep sequencing

#### Cleavage of target pool by IVTT-expressed I-MG11

All synthetic dsDNA were manufactured by Twist Bioscience (**Supplementary File S2**) at a scale of 1000 ng for each target. All samples were reconstituted in 50 μl nuclease-free water to make 20 ng/μl stock solutions. For pool cleavage, 2 μl (40ng) of each variant were mixed in a separate tube resulting in 0.22 ng/μl (4.3 x 10^8^ copies/μl) of each variant. IVTT reaction was prepared according to manufacturer’s instructions (PURExpress®, New England Biolabs) and 200 ng (2 μl of 100 ng/μl) of pET24a_IMG11 plasmid was added into 15 μl IVTT mix as the template for *in vitro* expression while no template plasmid was added in the control samples. After 2 hours of incubation at 37°C, 5 μl of pooled substrate oligos was added into the mix and incubated at 37°C. Samples were collected at different time point and used as PCR templates to amplify the non-cleaved remaining substrates using the Q5^®^ High-Fidelity DNA Polymerase (NEB) with the following primers: *Targets_FW: 5’_ttataggctgctggatgag*; *Target_Rv: 5’_ tcgccacctctgacttgag*.

#### NGS Processing and Quality Control

Raw paired-end fastq.gz files were processed by fastp v0.23.2 with a minimum quality score threshold of 30, and PEAR was used to merge overlapping reads. Probe sequences flanking the target region were used to extract the intervening relevant sequence using fuzzy matching (allowing up to 1 mismatch). Only sequences matching the target length of 30 bp were used for analysis.

#### Base Counting and Wild-Type Determination

For each sample, the pipeline counted the frequency of each base (A, C, G, T) at each position in the target region. The wild-type sequence was automatically detected from the control sample as the consensus sequence (most frequent base at each position). Reads with more than one mutation from wild-type (likely containing sequencing artifacts) were filtered out using the limit_to_hamming_1 parameter.

##### Analysis 1 (Figure 2a): Mutant-Only Specificity for identifying recognition window

Each mutant base was compared relative to the other mutant bases at the same position (the wild-type was excluded from this analysis) using the formula:

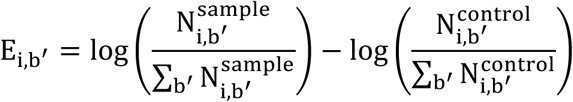

Where *E_i,b’_* is the enrichment of base *b’* at position *i*, and *b’* is the set of non-wild-type bases at position *I*, and 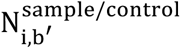 is the counts of base *b’* at position *i* in the sample and control, respectively. The resulting preference matrix was then row-normalized to create probability distributions among the three mutant bases, summing to 1.

##### Analysis 2 (Figure 2c): Global Wild-Type Comparison

Each base at each position was compared to the global wild-type fraction, resulting in enrichments of each base at each position that correspond to the relative cleavage of the oligo they represent using the formula:

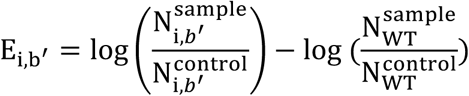

Where *E_i,b’_* is the enrichment of base *b’* at position *i*, and *b’* is the set of non-wild-type bases at position *I*, and 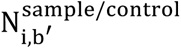 is the counts of base *b’* at position *i* in the sample and control, respectively, and 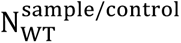 is the counts of full-length wild-type in the sample and control, respectively. To build the preference matrix, wild-type base values were set to the global wild-type enrichment difference log 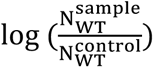, and mutant base values were set to E_i,b’_. Logo plots were then generated by exponentiating and row-normalizing the preference matrix.

##### Code availability

The NuCSOI pipeline handling the end-to-end pipeline is available as a modular command-line tool at https://github.com/Matt115A/NuCSOI and can be installed *via* pip. The bioinformatics pipeline used for target mutational scanning is available at https://github.com/Matt115A/nuclease-variant-profiler, along with install and usage instructions.

### Cleavage of targets by in vitro-expressed I-MG11

For assaying target DNA cleavage via in vitro-expressed I-MG11 enzyme, the enzyme was first expressed with PURExpress (NEB) for 2h at 37°C in a 20 µL reaction. Next, the expression lysate was incubated with 100-150 ng of substrate DNA for desired timepoint at 37°C. Prior to running samples on agarose gel, firstly RNase A (at final concentration of 50 μg/ml, QIAGEN) treatment was carried out for 10 min at room temperature, and then Thermolabile Proteinase K (at final concentration of 50 μg/ml, NEB) treatment for 30 min at 55°C was performed.

### Cleavage of targets by pure I-MG11

For cleavage of oligo variants, 40 ng (2 μl of 20 ng/μl) of each substrate were mixed in separate reaction tubes with 8 μl of enzyme (0.02 mg/ml) in the assay buffer. The reaction was stopped by collecting 2 μl samples into 2 μl of EDTA (50 mM) at 30 minutes (T1) and 60 min (T2). One microliter of collected samples was mixed with 3 μl sample buffer and analyzed using D1000 ScreenTape by a 4200 TapeStation Systems, according to the manufacturer’s instructions (Agilent).

For cleavage of supercoiled DNA, 5 nM of plasmid was used as substrate for 200 nM enzyme, and assay was stopped by collecting 2.5 μl of samples into 10 μl EDTA (50 mM) followed by visualization using 2% E-Gel™EX (Invitrogen) (see complete gel pictures at https://zenodo.org/records/18716102).

For the pH-rate profile, Tris-HCl buffer (50 mM Tris-HCl, 5 mM MgCl_2,_ 50 mM NaCl) was used for the pH range 7 to 9, and glycine-NaOH buffer (10 mM glycine-NaOH, 5 mM MgCl_2_) was used for assays at pH 10 to 12.

### Stability analysis of I-MG11 by nanoDSF

For conformational stability analysis by heat denaturation using nano differential scanning fluorimetry (nanoDSF), 10 µl of strep-tagged I-MG11 protein (0.62 mg/ml) loaded into capillaries, followed by heating, and simultaneously measuring intrinsic tryptophan fluorescence at 330/350 nm (for unfolding) them in a Prometheus Panta (NanoTemper) instrument. Double-stranded DNA was added at 50 µM final concentration. Target dsDNA and Non-Target dsDNA were prepared by annealing 30 bp complementary oligos as follows:

Target DNA 30 bp forward: TAAAAGGGTACGCGAGCTGGGTTCAAACCG
Target DNA 30 bp reverse: CGGTTTGAACCCAGCTCGCGTACCCTTTTA
Non-Target DNA 30 bp forward: GCGCGGGGATCTCATGCTGGAGTTCTTCGC
Non-Target DNA 30 bp reverse: GCGAAGAACTCCAGCATGAGATCCCCGCGC

### Structural Modelling

To obtain a structure prediction for the DNA/protein complex, two steps were performed: (1) the multiple sequence alignment (MSA) calculation and (2) the Boltz-1 calculations. The multiple sequence alignment (MSA) for I-MG11 was calculated with MMseqs2 (31). The Boltz-1 co-folding model (Wohlwend et al., 2025) utilizes evolutionary, structural, and physicochemical features analogous to methods implemented in AlphaFold Multimer (Evans et al., 2022) and AlphaFold3 (Abramson et al., 2024) and is capable of predicting of the complex of I-MG11 with the identified DNA recognition site and two Mg^+2^ ions important for catalytic activity. For the Boltz-a calculation, the diffusion samples were set to 25, the recycling steps to 10, while the predicted aligned error (PAE) and the predicted docking error (PDE) scores were extracted in the output models. The yaml input file for Boltz-1 is shown below.

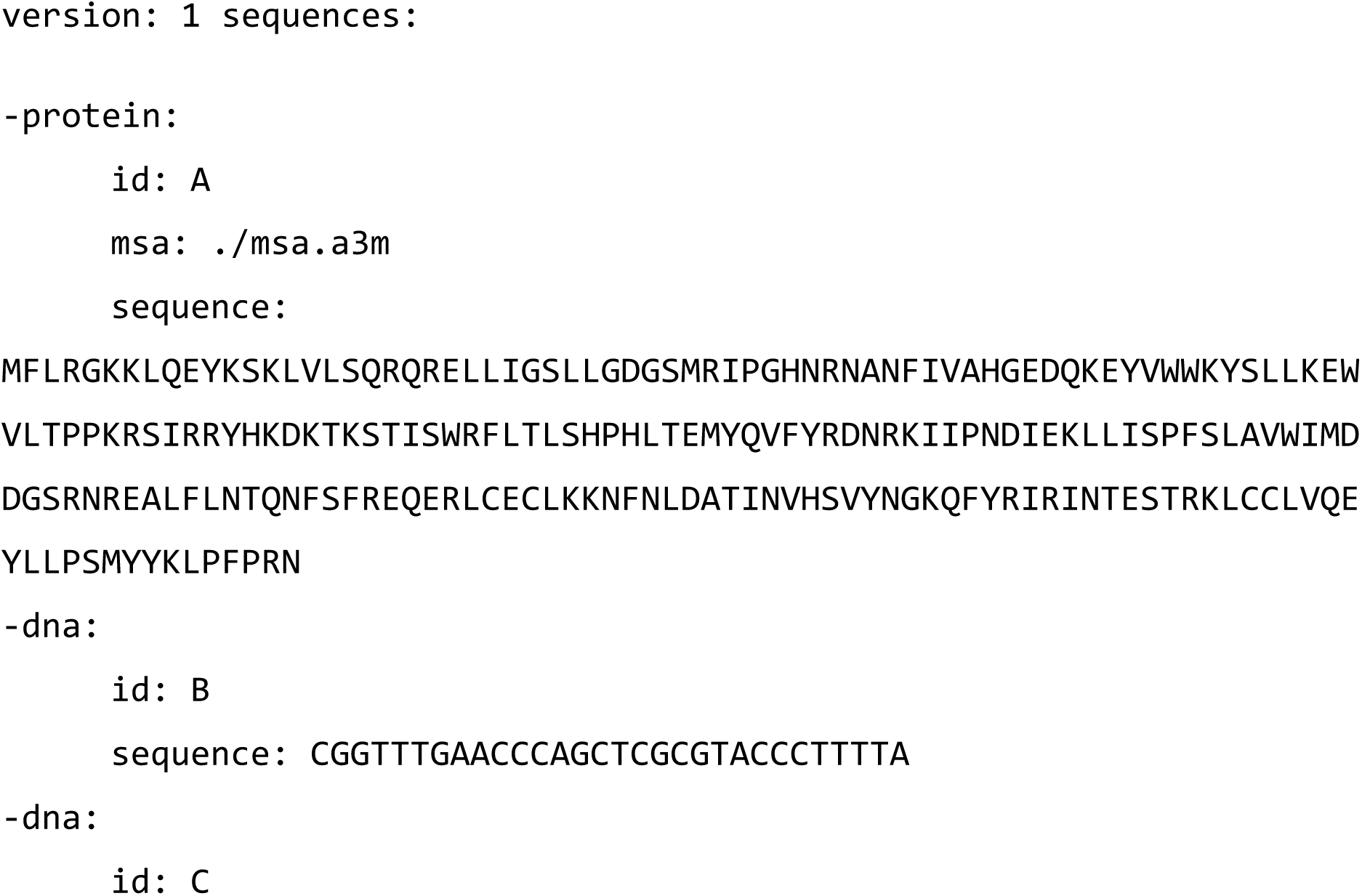
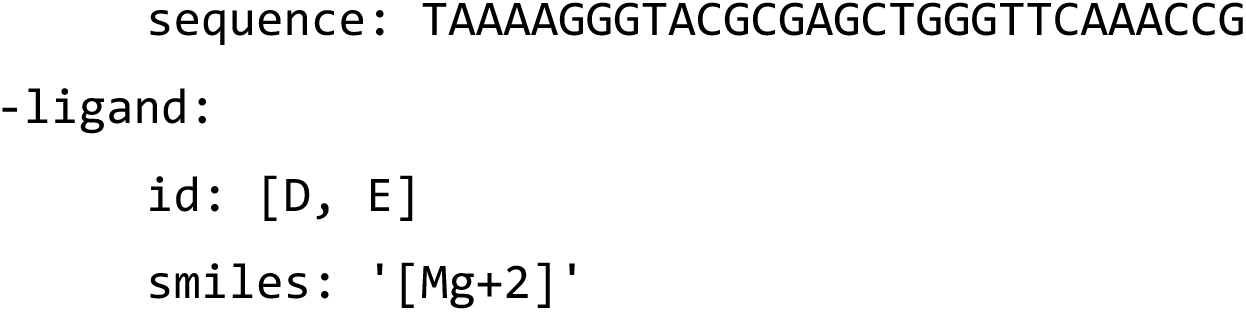

The visualization of the 3D models was performed with Maestro from Schrödinger (32).

## RESULTS

### Identification of meganuclease sequences from MGnify databas

We explored the sequence neighbourhood of the well characterized monomeric (single-chain) I-SceI meganuclease used as a bait and searched the non-redundant protein database using HMMER (23) resulting in ∼400 protein sequence predictions with significant homology to the query. The top 50 sequences (E-value <5.0e-14) were retrieved (**Supplementary Figure S1-A,B,C** and **FileS1**). Among these hits the two catalytic LAGLIDADG domains were highly conserved consistent with the query sequence (I-SceI) belonging to monomeric enzymes of this family (**Supplementary Figure S1-C**). Across a maximum likelihood phylogenetic tree based on the 50 hits, 16 variants were selected (**Supplementary Figure S1-B**, **Table S1**) and, after trying various cloning strains and strategies, 14 of them were successfully cloned into pET24 vector, under control of an IPTG-inducible T7 promoter. Despite using cells with a reduced basal expression (*E. coli* NEBExpress^®^ Iq cells, NEB), two of the selected variants did not yield clones with the correct gene sequence, possibly indicative a potentially toxic gene product (capable of genome cleavage). Such toxicity related to meganuclease expression in *E. coli* has also been reported previously (33, 34). The difficulties encountered during cloning, even in absence of induction of protein expression, suggested that achieving expression levels sufficient to purify and characterise meganucleases would be difficult. We opted to pursue a cell-free expression strategy to avoid time-consuming optimization steps and reduce the time from a t least 3-5 days to a few hours compared to traditional *in vivo* overexpression. We successfully expressed 13 out of the 14 variants (93%) using *in vitro* expression at the first attempt, showing the compatibility of this approach with meganucleases (**Supplementary Figure S1-D**). The resulting enzymes are readily produced via the cell-free expression and almost all are active (13 out of 14), in contrast to a previously reported approach that was only able to display 3 out of 6 meganuclease homologs on the surface of yeast (29). One of these, the 219 amino acid protein I-MG11 (Intron-encoded MGnify hit 11) was particularly interesting, as it showed only 28.3% pairwise identity to I-SceI (**Figure 1**). A protein BLAST of I-MG11 against the UniProtKB/Swiss-Prot database (https://www.uniprot.org/) (35) did not return any significant match, with the most similar protein having a highest subject identity of 31.96%. I-MG11’s metagenomic origin was located to an extreme hydrothermal marine environment from subseafloor sediment from Guaymas Basin (MGnify sample accession: SRS3033588). Next, the *E. coli*-codon optimized gene of the protein was cloned into pET24 vector for T7 polymerase-based *in vitro* as well as bacterial expression.

**Figure 1:**
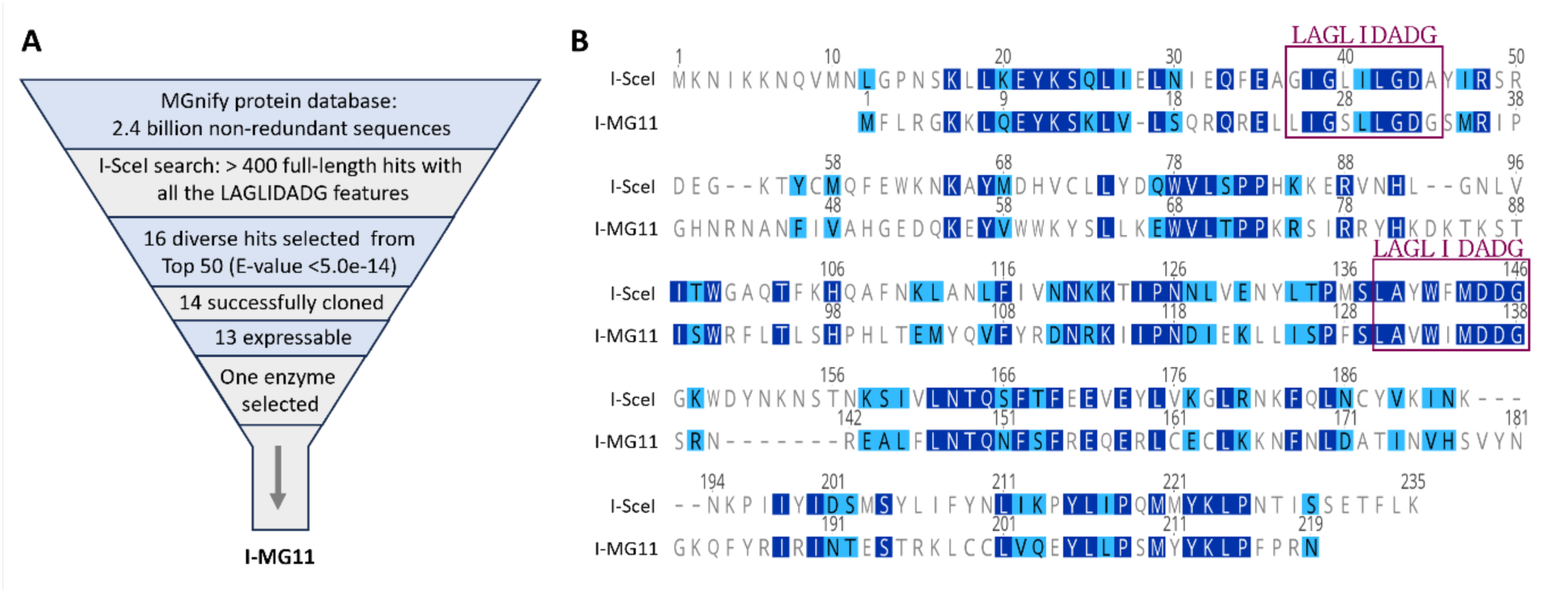
Metagenomic retrieval of potential meganuclease enzymes using I-SceI protein sequence as the query. **(A)** More than 400 protein sequences resulted in the initial search, from which, the top 50 sequences (E-value < 5.0e-14) were selected for further analysis. A high degree of conservation was observed within the catalytic LAGLIDADG domain across these sequences, confirming their classification in this family. Subsequently, From the successfully cloned and expressible variants, I-MG11, from a hydrothermal marine origin, was chosen for subsequent characterization. **(B)** Sequence alignment of I-MG11 and I-SceI shows 28.3% pairwise identity. Identical residues are highlighted in dark blue, and similar residues in light blue. Gaps in alignment are shown by a dash (-) symbol, representing insertion or deletion. Alignments were performed on Geneious Prime using Clustal Omega multiple alignments.

### Characterization of the recognition and cleavage site

The historical name of meganucleases, homing endonucleases (HEs), gives a clue to their target sequence specificity as they "home" into a particular DNA sequence within a genome, essentially facilitating the mobility and persistence of their own open reading frames. The genes encoding HEs are frequently integrated within self-splicing elements, such as Group I introns, Group II introns, and inteins (11, 12, 36).

To identify the recognition sequence of I-MG11, we thus inspected the original sequencing contig from the corresponding metagenomic samples on MGnify database (**Supplementary File S3**), which contained the genomic DNA sequence corresponding to I-MG11 protein. By searching the upstream and downstream region of the I-MG11 gene via a BLAST (Basic Local Alignment Search Tool) search, conserved exons of a 23s rRNA were identified in both up- and downstream regions. The putative cleavage site for an intron-encoded homing endonuclease would be reconstructed by fusing the two segments of the exon, which are split by the intron. Therefore, we removed the intron sequence surrounding the I-MG11, fused the sequences of flanking exons, and selected 15 base pairs from each exon in the fusion site to reconstruct a 30 bp putative target site of I-MG11 (**Figure 2-A**).

**Figure 2:**
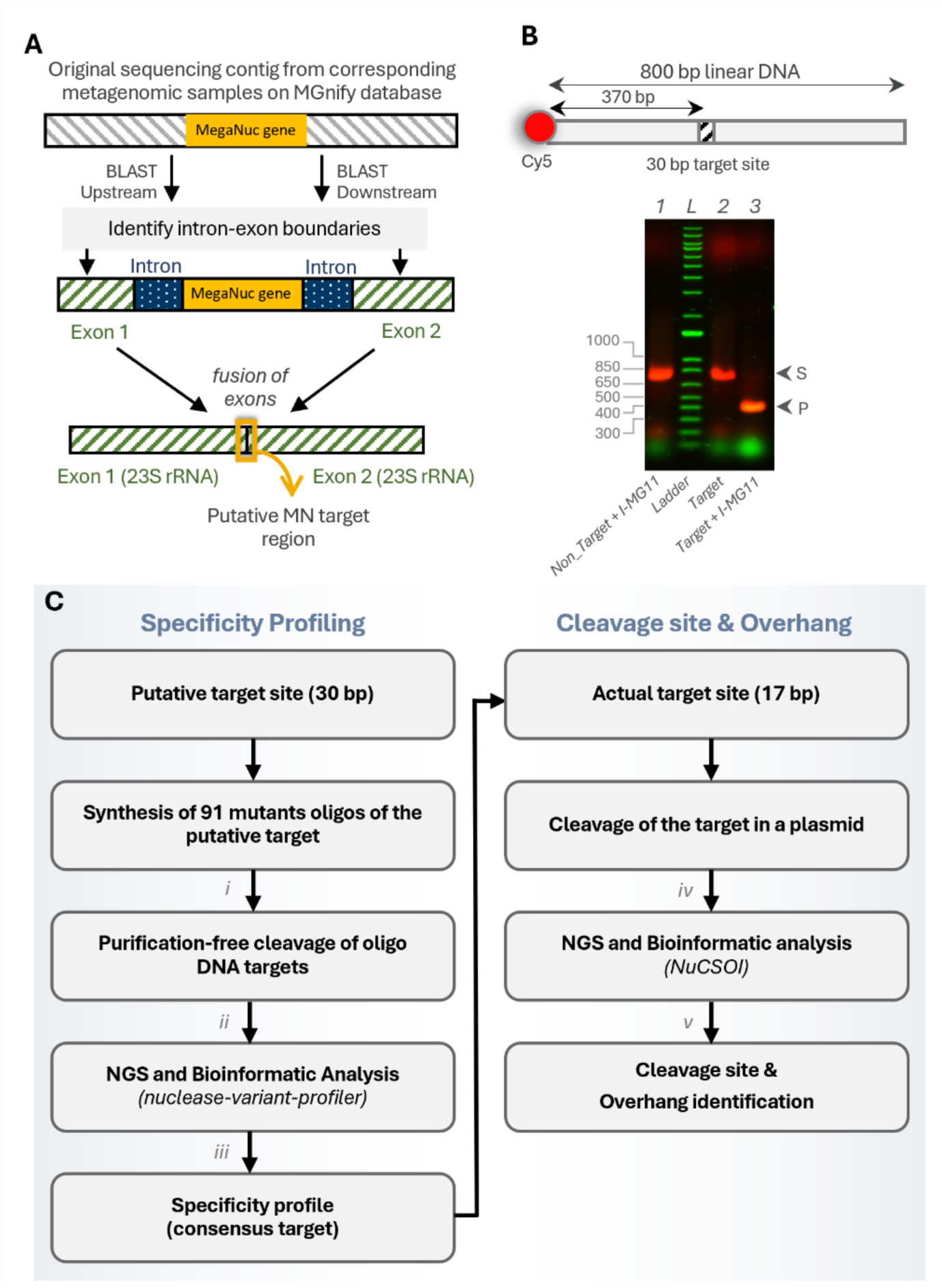
Identification of target site of intron encoded homing endonuclease I-MG11. **(A)** By inspection of metagenomic sequencing contig and identification of intron-exon boundaries via nucleotide BLAST and virtually removing the intron sequences by fusing the exon sequences, the putative cleavage site can be reconstructed. The putative 30 bp target of I-MG11 in the exon fusion site is cloned into a plasmid. **(B)** A linear dsDNA containing the HE cleavage site was amplified from the plasmid using Cy5-linked primers. Cleavage of the putative target is confirmed by running the samples on 1% agarose gel. *Lane 1*: Control1 as non-target DNA (I-SceI target site) treated with IVTT lysate expressing the I-MG11 enzyme; *Lane L*: 1 Kb Plus DNA Ladder (Invitrogen); *Lane 2*: Control2 as target DNA treated with only IVTT lysate (no-enzyme control). *Lane 3*: target DNA incubated with IVTT lysate expressing the I-MG11 enzyme for 1 hour. Fluorescence images of the Cy5 and Alexa Fluor 488 channels are shown in red and green, respectively. Bands for the substrate and product DNA fragments are indicated as S and P, respectively. **(C)** Workflow diagram for purification-free and NGS-based specificity profiling and cleavage pattern characterization. (*i*) Equimolar quantities of 91 oligos were pooled, representing the 30 bp putative sites mutated in each position to all remaining 3 alternative bases (90 variants) plus the wild-type parental sequence. (*ii*) PCR amplification of non-cleaved remaining oligos. (*iii*) A bioinformatic pipeline for specificity profiling based on NGS data is developed and applied. (*iv*) End-prep and adapter ligation. (*v*) The Nuclease Cleavage Site and Overhang Identification (NuCSOI) bioinformatic pipeline is developed and used for mapping the reads for the cleavage site analysis.

To increase the speed of discovery by obviating the need to heterologous expression and purification of the predicted protein, we designed a cleavage assay compatible with cell-free expression of meganucleases. We cloned the 30 bp putative target site in a plasmid and generated 800 bp linear DNA target by PCR amplification of a fragment containing the 30 bp target site. To directly monitor endounclease activity in the cell-free lysate, and to increase sensitivity in gel-based assays (in case of very low activities), we fluorescently labelled the target using a primer bearing a Cy5 (Cyanine5) fluorophore (**Figure 2-B**). The meganuclease was then expressed *in vitro* via an *in vitro* transcription–translation (IVTT) system (for 2 hours), and the lysate was incubated with the linear Cy5-labelled amplicon (for 1 hour). As controls, the IVTT mix without expression plasmid (i.e. lacking enzyme) and a non-target amplicon that contained I-SceI site but not the putative 30 bp site of the I-MG11 (no substrate control) were run in parallel. The nuclease reaction was stopped by addition of EDTA, based on separate confirmation that EDTA can stop the cleavage reaction and that Magnesium (Mg^2+^) is an activity-conferring metal ion cofactor for DNA cleavage (Figure S2). Fluorescent imaging of the samples run on agarose gel suggested cleavage of the amplicon containing the 30 bp putative site by the *in vitro*-expressed I-MG11 enzyme, while no cleavage of the amplicon containing I-SceI target site was observed (**Figure 2-B**). Next, we investigated the specificity of I-MG11 for its putative 30 bp binding region and attempted to identify the actual recognition and cleavage site embedded in this sequence.

### Deciphering the specificity profile of meganuclease I-MG11 using deep sequencing

In order to investigate the specificity profile of I-MG11 rapidly, we deployed a purification-free approach *via in vitro* expression of the enzymes, combined with deep sequencing of a target library, inspired by the approach of Thyme et al (37). Briefly, the 30 bp putative site was mutated at each position to all the remaining 3 nucleotides, resulting in a pool of wild-type sequence and single mutants (91 variants in total). All variants were individually synthesised as 499 bp double stranded DNA fragments, containing the 30 bp target site in the middle, flanked by non-target sequences (**Supplementary Figure S3**). All 91 oligos were pooled in an equimolar ratio and added into *in vitro*-expressed I-MG11 enzyme. Samples were collected at different time points and the nuclease enzyme reaction was stopped by addition of EDTA. The purified DNA samples were used as PCR templates for amplification of non-cleaved DNA fragments, followed by paired-end Illumina sequencing. After quality filtering of the deep sequencing data (see Methods & Materials), the frequency of each variant was determined from the NGS reads and compared to the control (IVTT mix with insertless plasmid template was used as a negative control) sample: the relative extent of cleavage for each variant during the incubation was quantified by normalizing the frequency of each variant in the output by its corresponding input frequency. In this analysis a higher score for the extent of cleavage corresponds to a decrease of abundance (de-enrichment) of particular variants in the output compared to the input. Background activity without added enzyme was tested for the longest timepoint (T4): this control showed no de-enrichment of specific oligos compared to the enzyme-treated samples (**Supplementary Figure S4-A, -B**), indicating a very low background of noise. Reproducibility was assessed using replicates of the enzyme-treated T4 sample – these are highly correlated (R^2^ = 0.99, **Supplementary FigureS4-C**).

To identify the recognition site of the nuclease, we assessed the relative tolerance for mutations at each of the 30 positions (**Figure 3-A**). The relative cleavage of mutants at each site indicates that selectivity is present between (and including) positions 8-24 (17bp) (Figure 3-a). To confirm this purification-free and NGS-based approach, we then compared the cleavage of 20 individual variants in the central region by the purified I-MG11 enzyme and verified the cleavage by electrophoresis (**Figure 3-B**). Both of the NGS-derived and gel-based cleavage patterns establish the same 17 bp as the recognition site of the enzyme (**Figure 3-C**). The cleavage of target variants by pure enzyme and the NGS-derived preference by in vitro-expressed lysate show a strong correlation (with Pearson’s correlation coefficients of R=0.774 for T1, and R=0.806 for T2 timepoints of enzymatic cleavage. **Supplementary Figure S5**), confirming the efficiency of the NGS-based approach.

**Figure 3:**
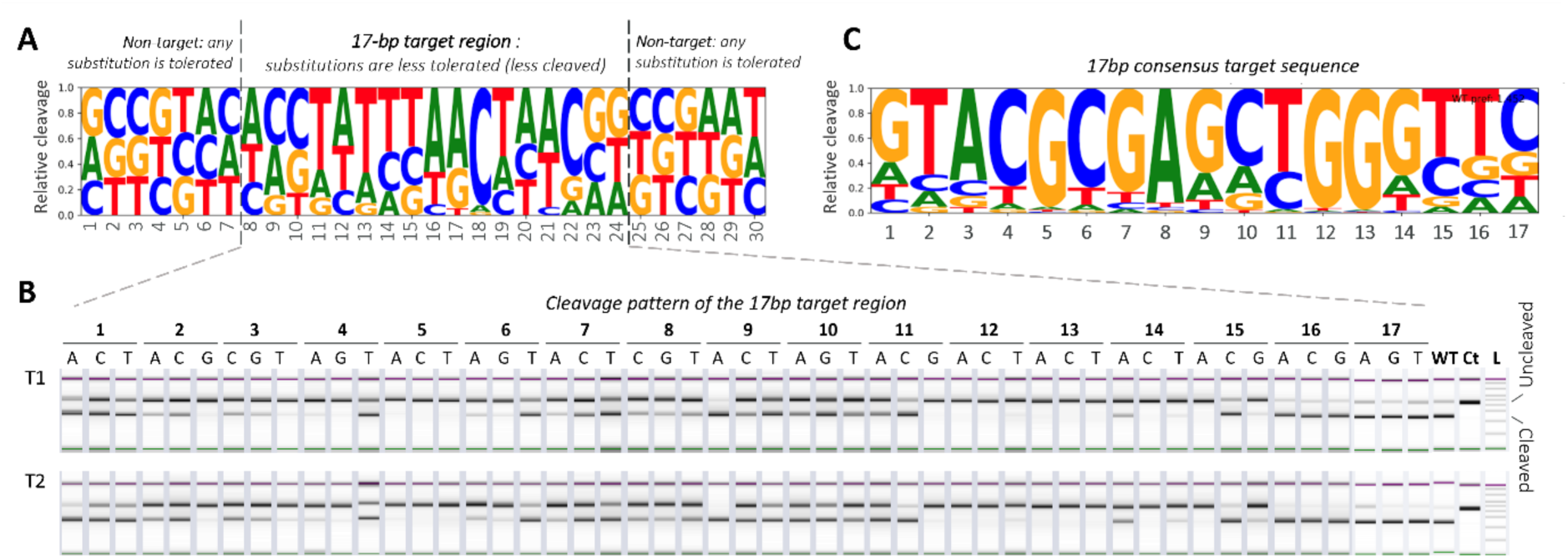
Cleavage specificity profiling of I-MG11 derived from deep sequencing. **(A)** Site-wise relative tolerance of the nuclease for mutations at each position. After incubation of the 91 variants of the target DNA (pooled at equimolar ratio) with IVTT-expressed lysate of I-MG11 enzyme for 8 hours, the uncleaved targets were amplified by PCR and subjected to NGS. Using NGS data, de-enrichment of each mutant base was compared relative to the other mutant bases at the same position. Logo represents preference matrix normalized to create probability distributions among the three mutant bases, summing to 1. Within the 30bp putative target, preference of the 17bp recognition sequence of the enzyme is clearly distinguishable. **(B)** Confirmation of NGS-based substrate scoping by cleavage of individual variants of the 17bp window by pure I-MG11 enzyme, and automated electrophoresis using TapeStation system (Agilent). Reaction is stopped by addition of EDTA at 30 minutes (T1) and 60 min (T2). WT: wild type target site; Ct, control reaction without enzyme; L: D1000 Ladder (Agilent). **(C)** Logo of the cleavage frequency of the 17 bp target window.

### The first monomeric meganuclease with palindromic four base pair 3’ overhangs

To identify the position and cleavage pattern (blunt or staggered) by I-MG11, we designed a pooled approach that quantifies reaction products based on paired-end next generation sequencing (NGS), dubbed *NGS-based Nuclease Cleavage Site and Overhang Identification (NuCSOI).* In this approach, a supercoiled plasmid containing the 30 bp target region was treated with IVTT-expressed I-MG11, followed by DNA clean up and enzymatic removal of overhangs prior to adapter ligation and paired-end sequencing (**Figure 4-A**). Blunt-end ligation of NGS adapters requires enzymatic removal of overhangs, a reaction that can be exploited to uncover the location of the overhangs by paired-end sequencing followed by single-base resolution cleavage site mapping by alignment against the reference (**Figure 4-B**). The results agreed with findings from classical run-off sequencing (i.e. Sanger sequencing from each direction towards the cleavage site), confirming a staggered cleavage with 3**’**-ends and 4 bp palindromic overhangs (**Figure 4-C**) (i.e. the inability of two-sided run-off sequencing to cover all bases at the 5’-end of the template strand near the cleavage site suggests a 3’ overhang on the non-template strand).

**Figure 4:**
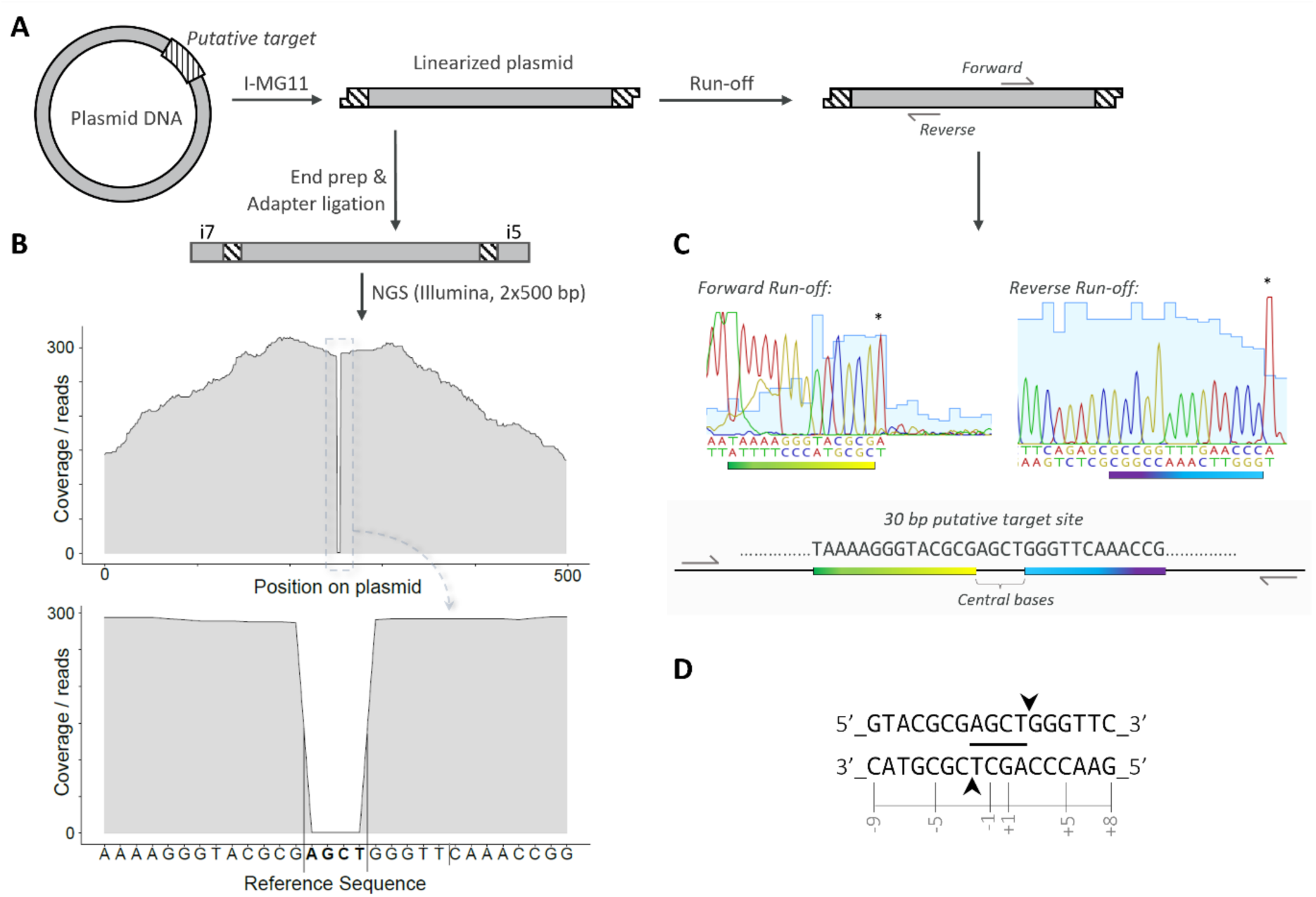
Identification of cleavage position and pattern by I-MG11 by NuCSOI (*NGS-based Nuclease Cleavage Site and Overhang Identification*) and run-off sequencing. **(A)** Supercoiled plasmid containing the 30 bp target region was cleaved by I-MG11 and the linearized plasmid was sequenced by NGS and then run-off sequencing to identify the cleavage site and pattern. **(B)** NGS of linearized plasmid includes enzymatic blunt ending prior to adapter ligation, which will remove any overhanging bases in the cleavage site. Each direction at the cleavage site is subjected to paired-end sequencing for single-base resolution cleavage site mapping. Position of cleavage and sequence of overhang is identifiable by alignment of reads to the respective circular plasmid reference, where abruption in local coverage indicates the cleavage site and the sequence of overhangs in each strand of DNA. Similar results are obtained by cleavage of supercoiled plasmid containing 20 bp central region (**Supplementary Figure S6**. **(C)** Run-off Sanger sequencing confirmation of central bases obtained from NGS approach. DNA sequencing chromatograms obtained by run-off Sanger sequencing are presented, and asterisks above peaks indicate a terminal adenine (A) nucleotide added by the template-independent activity of Taq DNA polymerase that was used for Sanger sequencing. **(D)** I-MG11 produces 4 bp staggered palindromic 3’ overhangs. Central (overhang) bases in the cleavage site of I-MG11 within the 17 bp actual target is obtained from NGS approach and the cleavage pattern is confirmed by run-off sequencing. The arrows point to the position where each strand is cleaved, forming 3’ cohesive ends indicated by underlined nucleobases (AGCT_3’) on either strand. Bases upstream of the central break are designated with negative numbers and bases downstream are given positive numbers, following the common convention for numbering of nuclease enzymes’ recognition site.

### I-MG11 is a thermostable enzyme that prefers high pH

To probe the versatility of I-MG11 to act on supercoiled plasmid DNA under different pH and temperature were tested. We first performed a pH scan (from pH 7 to 12), which established pH10 as the optimum pH for DNA cleavage activity (**Figure 5-A**). We then used this buffer and performed a temperature scan from 15 °C to 90 °C to assess the thermostability of the enzyme and identified the plateau between 55-65 °C as optimal for DNA cleavage activity (**Figure 5-B).** At these high temperatures and with excess enzyme (34, 38), 200 nM I-MG11 enzyme is able cleave 5 nM of supercoiled plasmid in 60 seconds at 60 °C, 22-times more than at 37 °C (**Figure 5-C**). The optimal temperature for most characterized meganucleases, such as I-SceI (34)(39), has been reported to be 37 °C, suggesting a unique role for I-MG11 for its ability to operate at high temperature.

**Figure 5:**
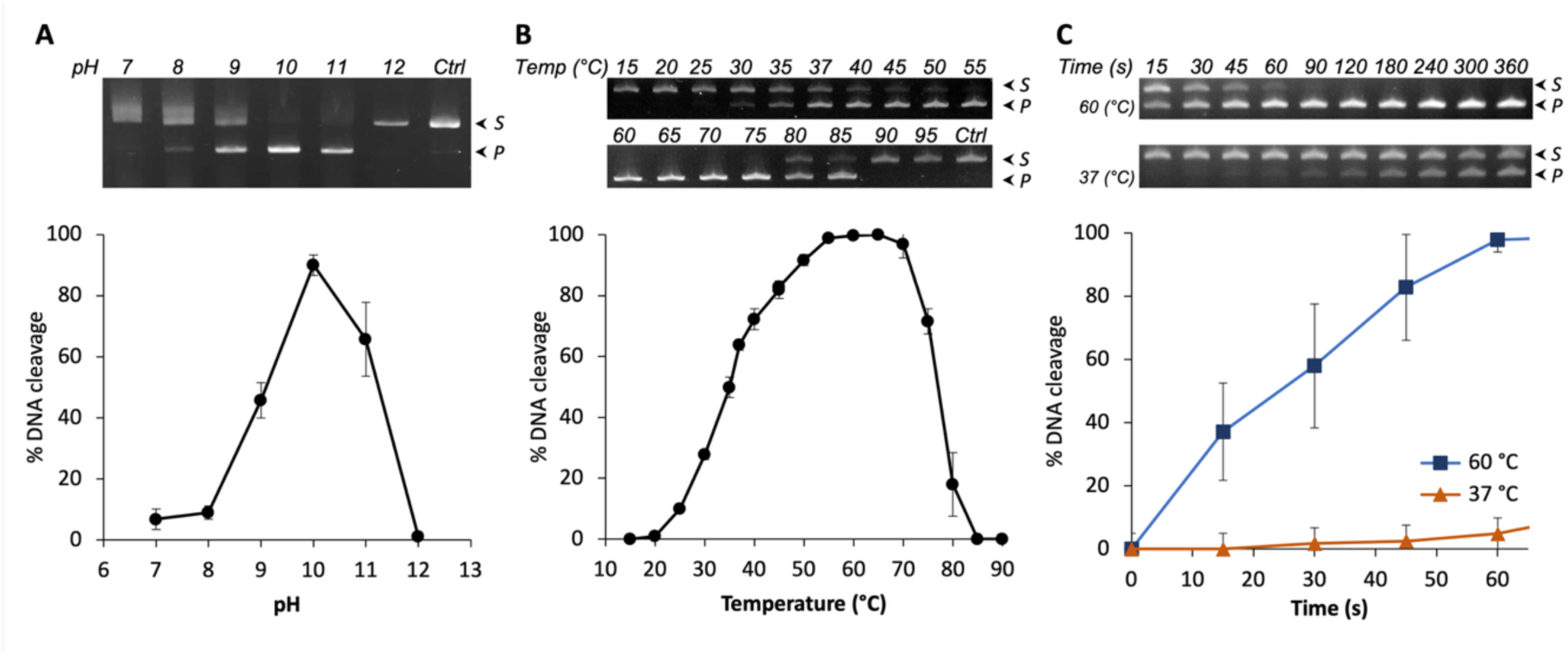
pH-rate dependence and temperature scan of I-MG11. **(A)** pH-rate profile of I-MG11 for DNA cleavage activity of supercoiled plasmid**. (B)** Effect of temperature on I-MG11 Activity. I-MG11 shows an activity plateau at 55-65 °C. **(C)** Supercoiled DNA (5 nM) cleavage over time of by purified I-MG11 meganuclease (200 nM) at 37 and 60°C. All the assays are performed in 10 mM Glycine-NaOH, 5 mM MgCl_2_, (pH 10.0) buffer. The values in all plots represent the percentages of cleavage read from corresponding bands by gel image analysis on 2% agarose gels (2% E-Gel™EX, Invitrogen). [Note that in the presence of EDTA in 2% gels, the cleaved linear DNA migrates faster than supercoiled DNA.] Substrate (S) and cleavage product (P) bands in each sample were quantified by ImageJ (**Supplementary File S4**). Error bars are the standard deviation of quantities obtained from 3 independent replicates. See https://zenodo.org/records/18716102 for pictures of complete gels.

**Figure 6A** shows Michaelis–Menten kinetics with *k_cat_* value of 0.76 min^-1^ and a *K_M_* of 72.7 nM (**Table 1** and **Supplementary Figure S7**), with mild substrate inhibition observed above 4 x *K_M_* (**Figure 6-B**). In order to probe the stoichiometry of reaction, sub-stoichiometric to excess concentrations of substrate were tested (**Figure 6** and **Supplementary Figure S8),** showing that the largest turnover is reached with 200 nM enzyme that was capable of cleaving more than 40 nM (2 µg) of supercoiled plasmid DNA in one minute (at [S] > *K_M_* is under optimal conditions: pH10, 60 °C).

**Figure 6:**
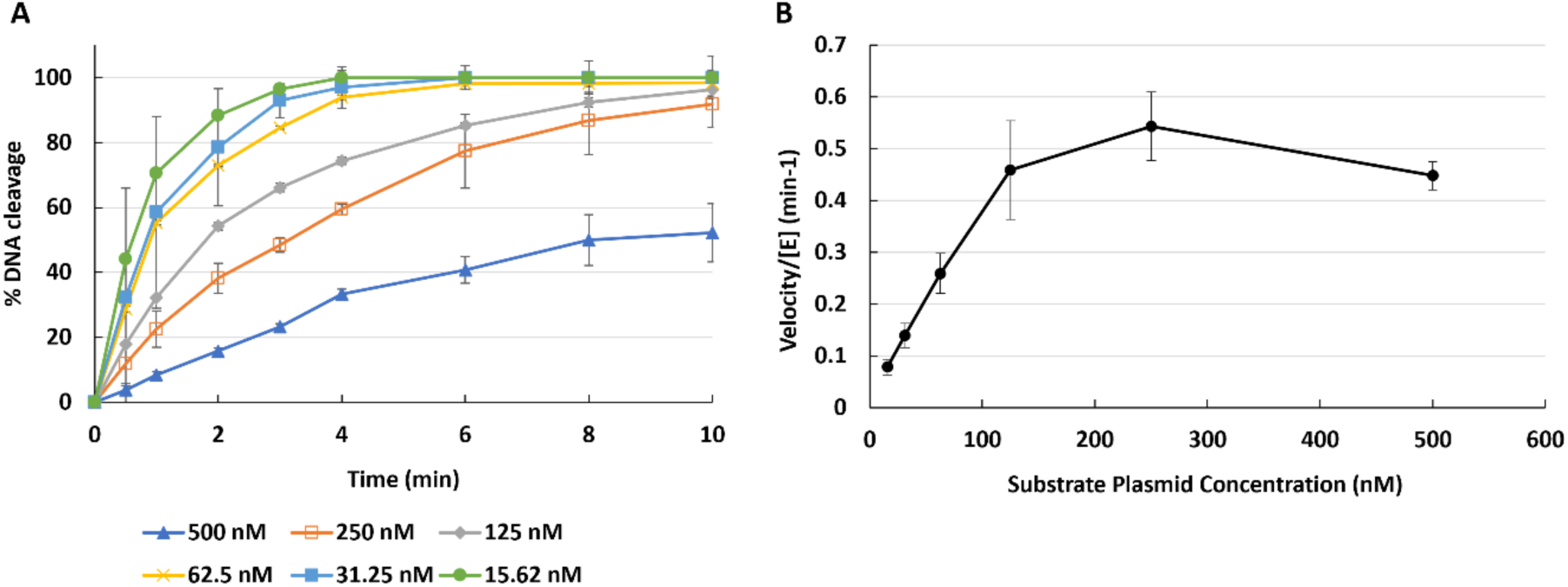
Cleavage kinetics of I-MG11. **(A)** Cleavage kinetics of the enzyme under different substrate concentrations. Supercoiled plasmid DNA at different concentration is treated with 200 nM of purified I-MG11 enzyme in at pH10 and 60°C. The values in the plots represent percentage of cleavage extracted from corresponding substrate and product band in each sample on 2% agarose gel images and quantified by ImageJ (**Supplementary File S4**). Error bars are the standard deviation of quantities obtained from at least 3 independent replicates. **(B)** Initial velocity of the enzyme against different substrate concentrations.

**Table 1:** Kinetics and stability analysis of the I-MG11 meganuclease. *k_cat_* and *K_M_* values were calculated by fitting the Michaelis–Menten equation to the experimental data (**Supplementary Figure S7**). Differential Scanning Fluorimetry (nanoDSF) analysis of the enzyme in absence and presence of 30 bp target or non-target DNA. Standard Deviations (S.D.) of at least three technical replicates are represented. Related nanoDSF output data are available in **Supplementary Figure S9**.

| $k_{cat}$ ($min^{-1}$) | $K_M$ (nM) | $T_m$ (°C) | | |
| --- | --- | --- | --- | --- |
|  |  | No DNA | + target DNA | + non-target DNA |
| **$0.76 \pm 0.05$** | $72.7 \pm 10.8$ | $74.2 \pm 1.3$ | $76.7 \pm 0.4$ | $68.5 \pm 0.22$ |

Finally, we assessed conformational stability of the pure Strep-tagged I-MG11 enzyme by nano Differential Scanning Fluorimetry (nanoDSF, **Table 1**), a label-free technique that measures changes in intrinsic protein fluorescence as temperature increases (39), giving a melting temperature (T_m_) of 74° C that increases with target DNA (to 77° C) and decreases with non-target DNA (to 69° C).

### Structural model of enzyme-DNA interactions

An atomistic model of the I-MG11 structure bound to the 30-bp DNA substrate containing the 17-bp recognition sequence, incorporating two Mg²⁺ ions essential for function, was generated using Boltz-1 (40), while AlphaFold 3 was unable to produce a DNA-bound model of I-MG11. Consequently, subsequent analyses focused on the Boltz-1–derived model. The top-ranked model exhibited a confidence score of 0.88, an interface predicted TM-score (ipTM) of 0.88, and a complex interface predicted docking error (iPDE) of 0.53. The model preserves monomeric LAGLIDADG-like motifs characteristic of I-SceI (41), including the catalytic aspartate residues (**Figure 7A**), but also displays substantial differences relative to I-SceI. Specifically, I-MG11 lacks an N-terminal helix due to an 11-amino-acid deletion; contains two-residue insertions between the β1 and β2 antiparallel strands that increase the accessibility of the intervening loop to the DNA major groove; includes a two-residue insertion between the β3 and β4 strands that slightly lengthens these strands; features a seven-residue deletion in the loop connecting β7 and β8, reducing its penetration into the DNA major groove; and harbors a five-residue insertion between β9 and β10 that extends these antiparallel β-strands. In addition, the DNA helix in the I-MG11 complex displays more pronounced bending, consistent with a stronger induced-fit upon binding (**Figure 7A**). The model also suggests specific protein–DNA interactions (**Figure 7B**): multiple positively charged residues in I-MG11 (R4, R36, R189, K74, K84) are oriented toward the DNA, consistent with electrostatic interactions with the negatively charged phosphate backbone, while polar residues (S139, N148) appear to form hydrogen bonds with DNA.

**Figure 7:**
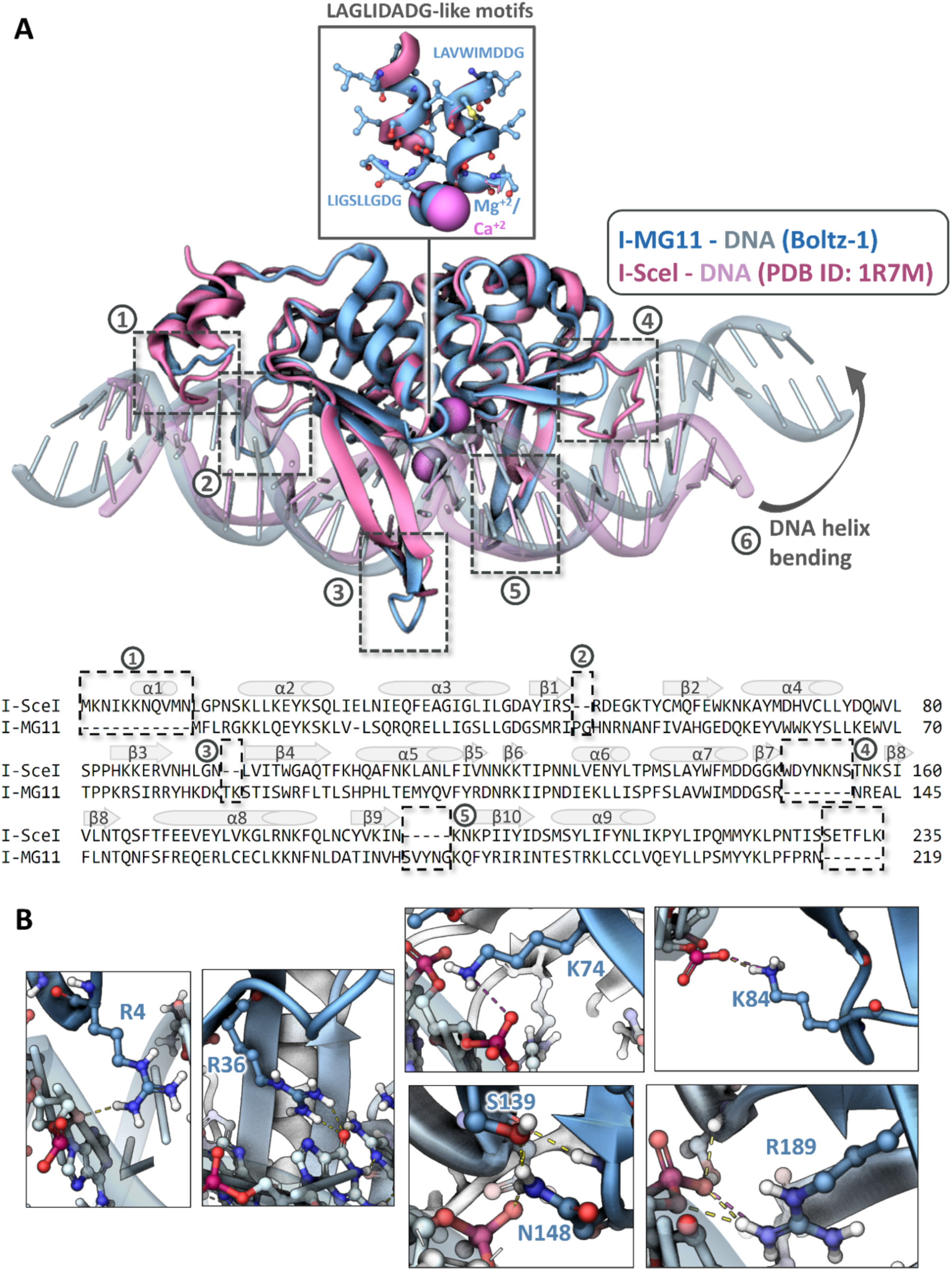
I-MG11/DNA co-folding model. **(A)** *Top*: structural alignment of the I-MG11/DNA Boltz-1 model (**Supplementary File S5**) with I-SceI meganuclease bound to DNA (PDB ID: 1R7M). The two meganucleases and the DNA helices are shown in cartoon representations in blue and pink colours for I-MG11 and I-Scel, respectively. The alignment of the LAGLIDADG-like motifs is shown in the figure inset and InDels that distinguish bound I-MG11 and I-Scel are highlighted. *Bottom*: amino acid sequence alignment of I-MG11 and I-SceI, where α helices and β sheets secondary structures are highlighted by cylinders and arrows, respectively. The differences between the two enzyme complexes reflect the InDels in detected on the structural overlay of the top panel. **(B)** Specific key interactions between I-MG11 residues (dark blue) and DNA bases (light blue).

To probe the role of InDels in functional divergence, we also generated Boltz-1 models of an InDel-free version of I-MG11 (I-MG11-InDel-free, **Supplementary File S6**), where all the InDels in I-MG11 relative to I-SceI protein sequence (**Figure 7B**) were reverted. This comparison creates an opportunity to distinguish the contributions of the diverging backgrounds of the two enzymes vs the effects of portable InDel features on substrate recognition. If the latter are functionally relevant we would expect their removal to be reflected in structural rearrangements around the substrate recognition site. Overall, the reversion of InDels is generating a I-MG11 more similar to I-SceI. Structural alignments of the I-MG11-InDel-free with I-SceI (**Supplementary Figure S10**) suggest that the reversion of InDels resulted in removal the extensions of β4, β9 and β10 sheets, as well as shortening the extended loop between β1 and β2 antiparallel sheets, causing a less closed DNA-binding pocket compared to I-MG11 (**Supplementary Figure S11).** Given that changes affect the DNA binding cleft of I-MG11, a role of the InDels in governing DNA substrate recognition, it is conceivable that essentially portable InDels influence the specificity profile of this meganuclease. In addition, a more compact DNA-binding pocket through InDels may also play a role in the I-MG11’s high thermostability, though this requires further investigation.

## DISCUSSION

I-MG11 is a single-chain monomeric LAGLIDADG meganuclease, which has no significant match in a sequence comparison against functionally annotated proteins in the UniProtKB/Swiss-Prot databases, suggesting that its functional characterisation breaks new ground, despite initially being derived from a homology comparison. A previous BLAST-search had sampled a narrower homology space (57% vs 28%) and achieved scaffold diversity but remained identical in the functional properties of the characterised enzymes (29). The depth of MGnify and the loosening of sequence homology constraints made broader database harvesting possible and increased the chances of identifying functionally distinct meganucleases. Experimental elucidation of I-MG11’s cleavage profile then showed that further sequence deviation from known enzymes lead to discovery of novel features. To this end, a library approach supported by next generation sequencing provided sufficient experimental depth to overcome the lack of predictive prowess for an enzyme with low homology to existing enzymes that is placed in entirely novel, unannotated sequence space. Specifically, I-MG11’s 17 bp recognition site has no significant similarity to previously characterized meganuclease enzymes (**Supplementary Figure S12** and **Table S2**). There is an immediate use for I-MG11’s recognition properties expanding the toolkit of highly specific molecular scissors. In addition, I-MG11 generates 4 base pair *palindromic* 3′ overhangs. However, the capacity to generate palindromic DNA overhangs has traditionally been associated with homodimeric LAGLIDADG homing endonucleases, such as I-CreI and I-CvuI, which bind palindromic or near-palindromic DNA target sequences (42–44). Whereas monomeric enzymes in this family, such as I-SceI, were long considered restricted to cleaving asymmetric, non-palindromic target sites, hence generating non-palindromic overhangs (42–44). To our knowledge, I-MG11 is the first monomeric meganuclease to generate such 3′-palindromic ends by cleaving a non-palindromic target site. Besides providing evolutionary and mechanistic insights, these properties may potentially be used in meganuclease-mediated recombinant DNA fragmentation and assembly (45, 46) to specifically generate overhangs suitable for universal adapters and thus may relax fragment-specific compatibility.

A notable feature of LAGLIDADG meganucleases is their high isoelectric point (pI). For the top 50 MGnify hits, the mean pI is unusually high around ∼10 (**Supplementary File S1**), suggesting that these enzymes carry a net positive charge under physiological, near-neutral conditions. Mapping of the charge distribution on a structure prediction of I-MG11 shows that positive surface patches are located in the DNA-binding pocket (**Supplementary Figure S13**). Positively charged surface patches are a common structural feature of DNA-binding proteins involved in promoting electrostatic interactions towards the negatively charged phosphate groups of the DNA backbone (47–50), and maintaining non-specific binding when sliding in search of the specific target site (47, 51, 52) via a random walk (53, 54). However, in *in vitro* assays, I-MG11 exhibits maximal activity around pH 10 (**Figure 5**), seemingly avoiding net positive charge (similar to other LAGLIDADG enzymes such as I-SceI and I-WcaI (40)), This may increase free diffusion and hopping along the DNA as has been shown by increase in salt concentration that weakens the electrostatic interactions (53–55).

MGnify’s metadata informs us about I-MG11’s origin in subseafloor sediments from the Guaymas Basin, explaining its thermostability – and demonstrating the value of sequencing projects from extreme environments. A thermostable nuclease enables genome editing in thermophiles (56, 57), is also compatible with steps that are carried out at elevated temperatures (e.g. in Loop-mediated isothermal amplification (LAMP) (58) and Gibson assembly (59)). Multi-step workflows can be conceived in which restriction/ligation steps are performed at lower temperatures, while steps carried out by a specific nuclease are triggered at high temperature.

In this discovery campaign, *in vitro* expression overcame the problem of inefficient heterologous expression of genes from unusual environments, where little or no ‘local’ biochemical characterisation was available (as in 13 of 14 cases tested in this work). Our NGS-based methods also employ cell-free expression, so that experimental discovery and elucidation of specificity and cleavage pattern (by NuCSOI: NGS-based Nuclease Cleavage Site and Overhang Identification) can be performed without host constraints. This approach thus provides access to unique biocatalysts offered by the billions of candidates in MGnify, dispensing with the need to isolate and culture the species from which it originated. A comparison of known and novel enzymes can provide mechanistic and functional insight, in this case highlighting the role of InDels as quasi portable features that impact substrate recognition. Given that only a miniscule fraction of open reading frames in MGnify has been experimentally annotated, providing *bona fide* functional annotation is crucial to unlock the full potential of this and other sequence databases for diverse biomedical and biotechnological applications.

## ACKNOWLEDGMENTS

We would like to thank Felix Langer of Dr. Robert Finn’s group (the MGnify DB developers, EMBL-EBI) for help in finding the original DNA sequencing contigs of the MGnify protein hits.

## FUNDING

The work was supported by AstraZeneca. M.P. and F.H. acknowledge support from the Horizon Europe programmes *BlueRemediomics* (101082304) and *BlueTools* (10058118) (implemented by Innovate UK) and L. B. a studentship from the EU H2020 Marie Curie Network MMBio (721613).

## DATA AVAILABILITY STATEMENT

The sequence of I-MG11 was deposited on ENA with accession number ERZ29115786. All next generation sequencing reads can be downloaded from the SRA (accession numbers PRJNA1426418 for NUCSOI and PRJNA1426412 for substrate profiling). The codes for cleavage site identification of nucleases by NUCSOI (10.5281/zenodo.18715192) and for interpretation of substrate profiles (10.5281/zenodo.18715192) were deposited in the Zenodo repository. Pipelines for NuCSOI (https://github.com/Matt115A/NuCSOI) and for target mutational scanning (https://github.com/Matt115A/nuclease-variant-profiler) are also available on GitHub along with install and usage instructions. Complete pictures of gels shown in this work can be found at Zenodo (https://zenodo.org/records/18716102). A list of the top 50 hits retrieved from MGnify is eclosed as Supplementary File S1, synthetic library dsDNA sequences in S2, the contig of the corresponding metagenomic samples in MGnify in S3, reaction product analysis in S4, the Boltz-1 I-MG11/DNA co-folding model in S5 and Boltz-1 models of an InDel-free version of I-MG11 in S6.

## AUTHOR CONTRIBUTIONS

G.M.D., conceived the project, conducted all experiments and prepared the original draft. M. Pa. wrote code and carried out NGS analysis. L. B. contributed to optimization of experiments and project design. C.A. and J.C.M. performed AlphaFold modelling. C.W., M. P., and F.H. supervised the project. All authors contributed to writing the manuscript.

## CONFLICT OF INTEREST

G.M.D., C. A., J. C. M. C.W. and M. Pa. are (or were) employees of AstraZeneca. The other authors declare no competing interests.

