## Supplementary Information for "A metagenomic thermostable monomeric meganuclease with novel specificity and unique palindromic 3’ overhangs"

### for

### 1. Supplementary Figures

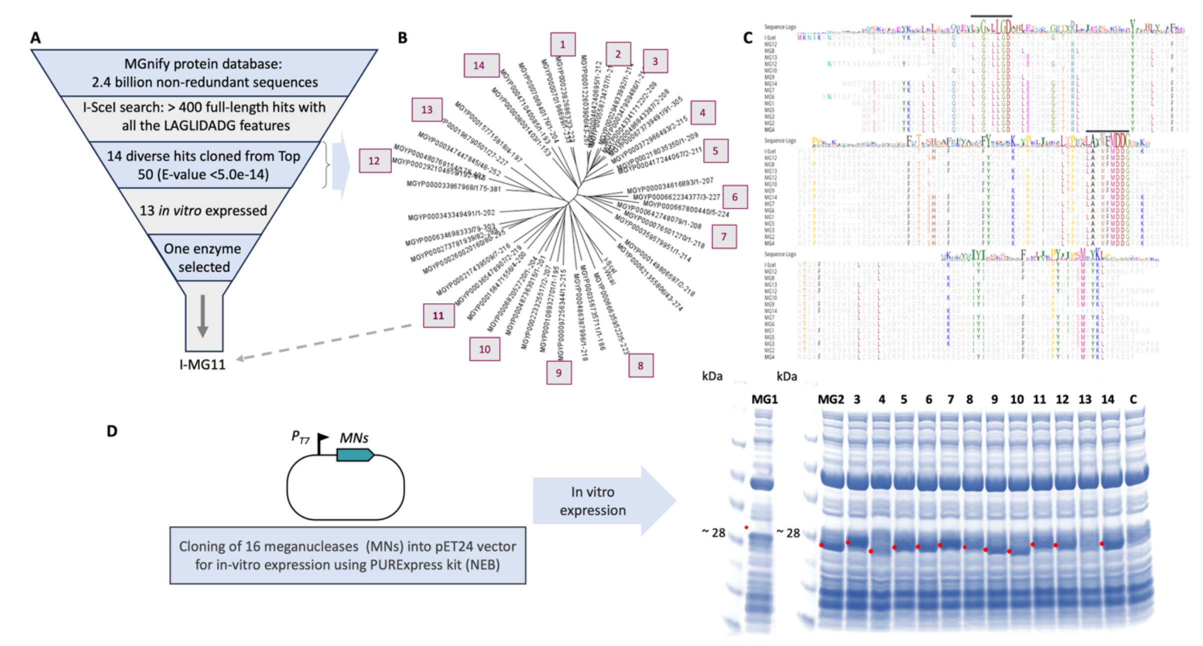

**Figure S1:** Retrieval of putative LAGLIDADG meganucleases from the MGnify database using the sequence of the enzyme I-SceI as the query. **(A)** More than 400 protein sequences resulted in the initial search, from which, the top 50 sequences (E-value < 5.0e-14) were selected for further analysis. A high degree of conservation was observed within the catalytic LAGLIDADG domain across these sequences, confirming their classification within the LAGLIDADG enzyme family. Subsequently, 14 variants were randomly selected from a maximum likelihood phylogenetic tree constructed using the 50 sequences. Among them I-MG11 (MGnify protein identifier: MGYP000365478907), from extremely hot subseafloor sediment of Guaymas Basin (MGnify sample accession: SRS3033588), which had no significant match in protein BLAST against UniProtKB/Swiss-Prot database (highest subject identity of 32%, with E value of 8e-26), was chosen for subsequent characterization. **(B)** Maximum likelihood phylogenetic tree of the top 50 hits. **(C)** Sequence alignment of the 14 randomly selected proteins that shows conservation of the LAGLIDADG catalytic motif (indicated by a line on top). **(D)** Initially, 16 proteins were codon optimized for *E. coli* expression, ordered as synthetic genes, and cloned into pET vector for T7-based expression. Cloning in *E. coli* NEBExpress® Iq Competent cells was successful for 14 of the enzymes except for two of them: MGYP000621355806 and

MGYP000273781939. *E. coli* NEBExpress® Iq reduces the basal expression, hence overcoming associated toxicity of the gene of interest in *E. coli*. Despite several repeats, cloning of these two genes in different hosts has failed, suggesting a high toxicity of these enzyme, probably due to high activity of these enzymes against *E. coli* chromosome. In vitro expression of the cloned enzymes using the PURExpress kit was successful, except for MG15, after a 4h expression according to manufacturer's instructions.

| EDTA | + | + | + | + |
|---|---|---|---|---|
| **$Mg^+$** | - | + | - | - |
| **$K^+$** | - | - | + | - |
| **$Zn^+$** | - | - | - | + |

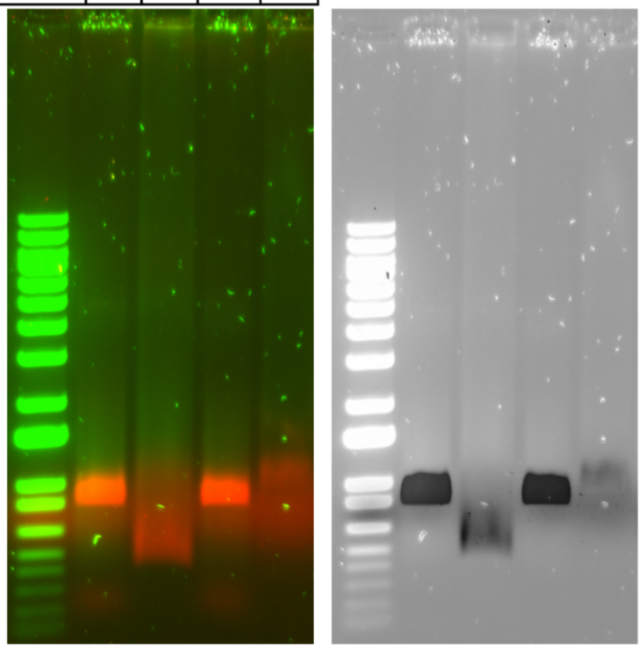

**Figure S2**: Confirmation of Magnesium ($Mg^{2+}$) as the catalytic ion for I-MG11 activity. The reaction of I-MG11 (0.02 mg/ml) with fluorophore-linked DNA (250 nM) target was quenched by addition of excess EDTA (to a final concentration of 75 mM), then the sample was divided into 4 tubes and 100 mM of either potassium chloride, magnesium acetate or zinc acetate was added and the mixture incubated for 4 hours. In the left panel the fluorescence imaging of the agarose gel is presented: green represents the Alexa Fluor 488nm channel, red represents Cy5 channel. In the right panel, the inverted image is presented for ease of readability.

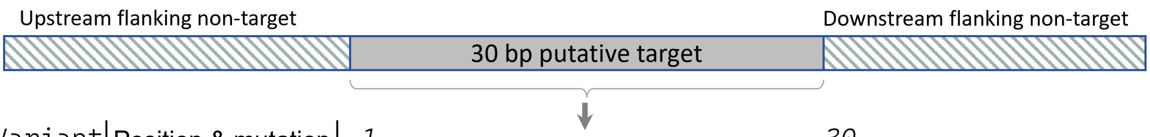

| Variant | Position & mutation | 1............................30 |
|---|---|---|
| 1 | parental | **TAAAAGGGTACGCGAGCTGGGTTCAAACCG** |
| 2 | 1A | A............................. |
| 3 | 1C | C............................. |
| 4 | 1G | G............................. |
| 5 | 2C | .C............................ |
| 6 | 2G | .G............................ |
| 7 | 2T | .T............................ |
| 8 | 3C | ..C........................... |
| 9 | 3G | ..G........................... |
| 10 | 3T | ..T........................... |
| . | . | . |
| . | . | . |
| . | . | . |
| . | . | . |
| . | . | . |
| 83 | 28A | ...........................A.. |
| 84 | 28G | ...........................G.. |
| 85 | 28T | ...........................T.. |
| 86 | 29A | ............................A. |
| 87 | 29G | ............................G. |
| 88 | 29T | ............................T. |
| 89 | 30A | .............................A |
| 90 | 30C | .............................C |
| 91 | 30T | .............................T |

**Figure S3:** Variants of synthetic double-strand DNA for all single mutants of the putative 30 bp target site. Together with the parental target, total 91 variants were designed and commercial oligonucleotide synthesis was carried out by Twist Biosciences.

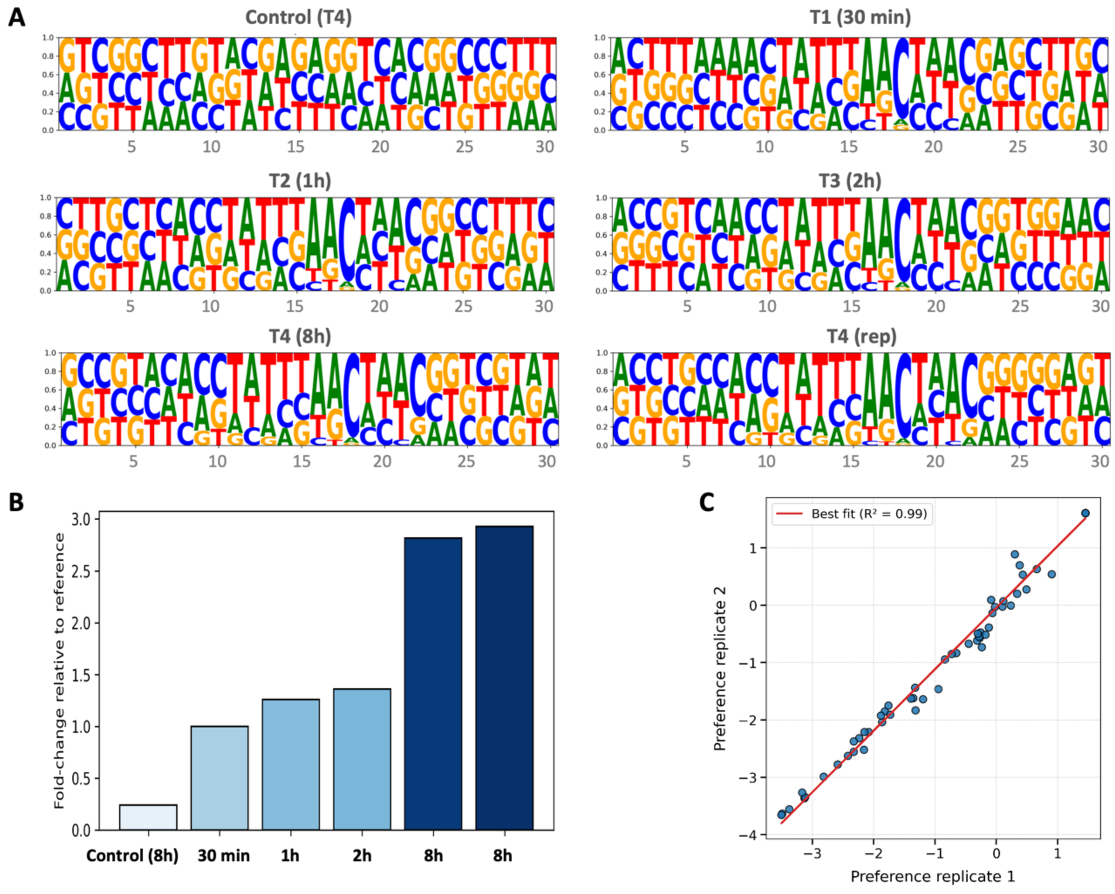

**Figure S4:** Deep sequencing analysis of pooled target library after treating with in vitro-expressed I-MG11 enzyme across different timepoints. **(A)** Representation of the relative tolerance for mutations at each position catalysed by the I-MG11 nuclease. After incubation of the 91 variants of the target DNA (pooled at equimolar ratio) with IVTT-expressed lysate of I-MG11 enzyme for 0.5, 1, 2 and 8 hours, the uncleaved targets were amplified by PCR and subjected to NGS. Using abundance obtained from NGS data, de-enrichment of each mutant base was compared relative to the other bases at the same position. Logo represents preference matrix normalized to create probability distributions among the three mutant bases, summing to 1. **(B)** Fold change overtime in the pool relative to starting time (T0 reference). **(C)** The longest timepoint (T4=8h) was run in two replicates to ensure reproducibility. Replicates of the enzyme-treated T4 sample are highly correlated.

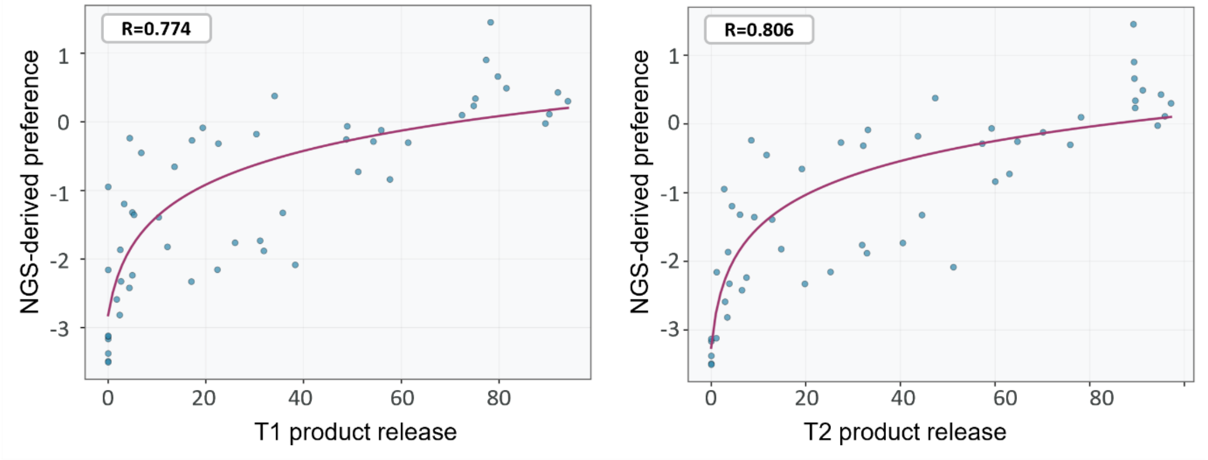

**Figure S5:** Relationship between the cleavage of target DNA variants by pure enzyme, and the NGS-derived preference by *in vitro*-expressed I-MG11. Pearson's correlation coefficients (R) are indicated.

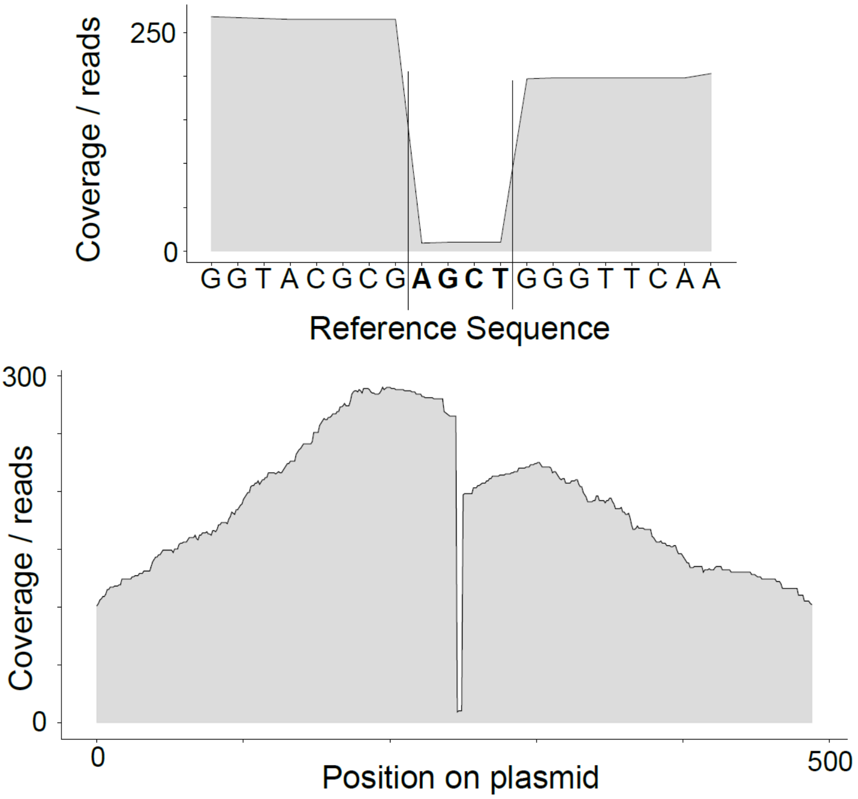

**Figure S6:** Single-base resolution cleavage site and overhang mapping on the 20 bp central bases as substrate in a supercoiled plasmid. Alignment of reads to the respective circular plasmid reference, identifies cleavage site and overhang sequence by abruption in local coverage.

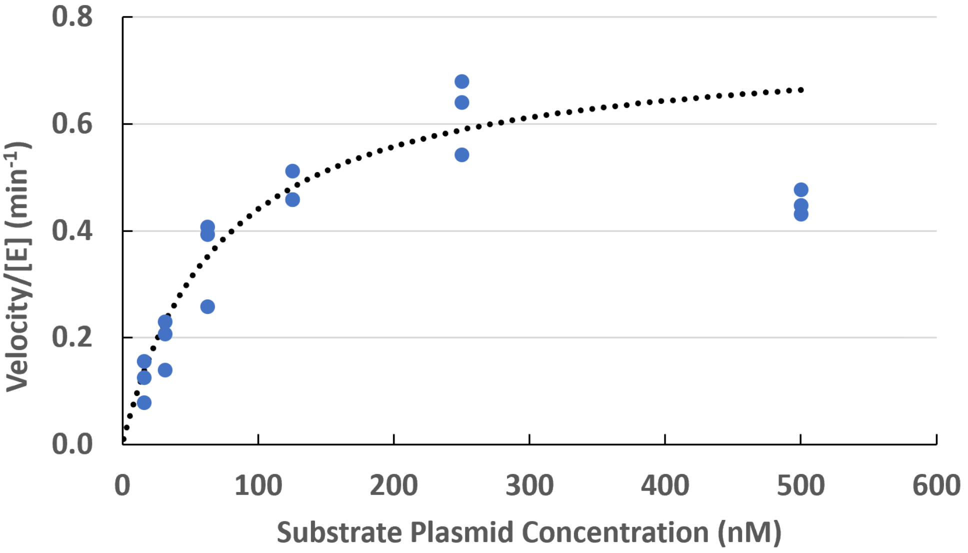

**Figure S7**: Kinetic analysis of the enzyme I-MG11. Initial reaction velocities were measured at varying concentrations (0 to 500 nM) of the supercoiled plasmid substrate in 10 mM glycine-NaOH, 5 mM $MgCl_2$, pH 10. The dotted line represents the nonlinear fit of the experimental data (solid circles) to the Michaelis-Menten equation. At each concentration of the plasmid substrate at least three data points were collected.

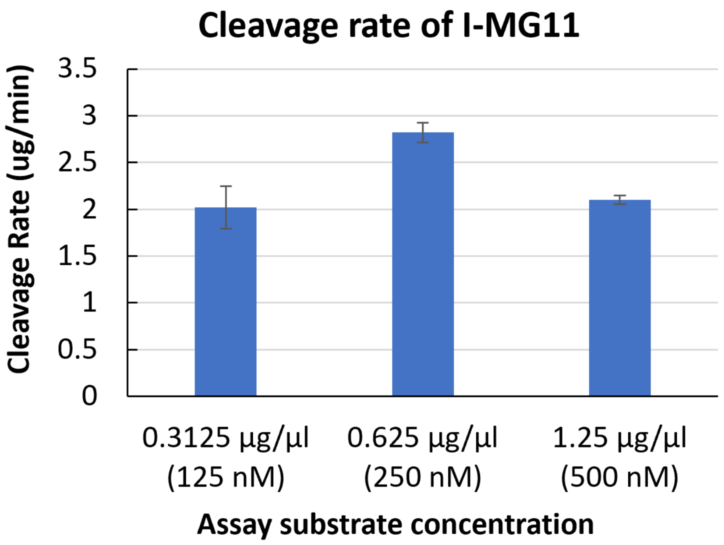

**Figure S8**: Maximal cleavage rate of I-MG11. Under excess substrate conditions (e.g. higher than $K_M$ = 73 nM), I-MG11 meganuclease (200 nM) cleaves more than 2 micrograms of supercoiled plasmid substrate in a minute under optimized assay condition of 10 mM Glycine-NaOH, 5 mM $MgCl_2$ buffer (pH10) and 60 °C.

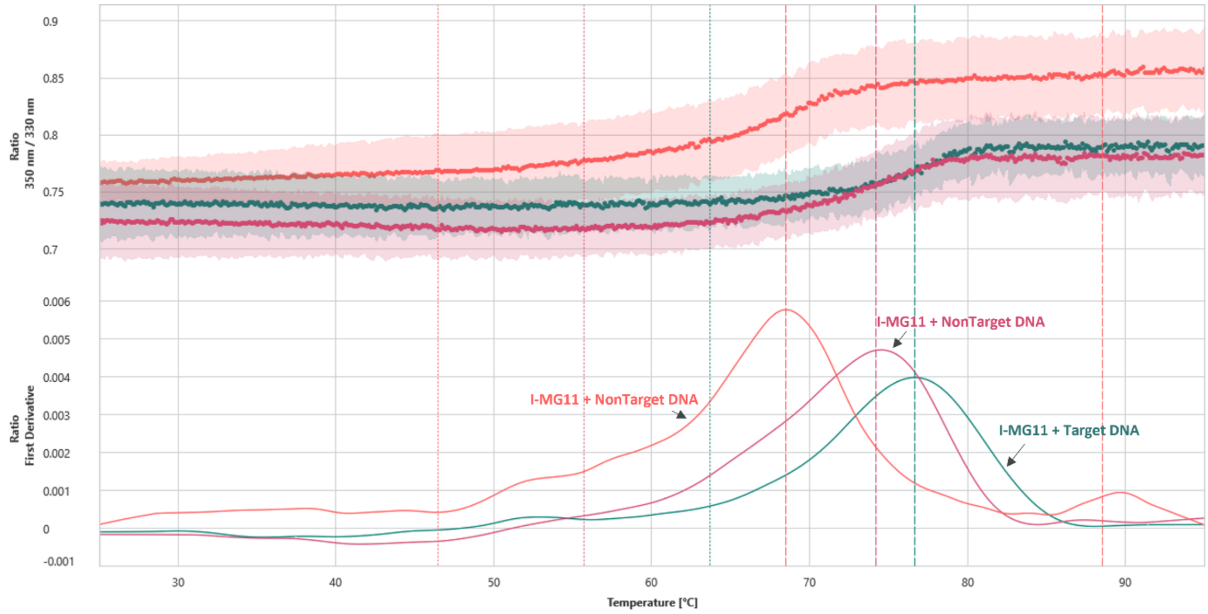

**Figure S9.** Thermal unfolding analysis by nano Differential Scanning Fluorimetry (nanoDSF) obtained from measurements with a NanoTemper Prometheus Panta instrument (NanoTemper®). *Top:* ratio of emission at 350 nm and 330 nm against changes in temperature. *Bottom:* first derivative of the ratio (d(F350/F330)/dT) used to identify the inflection point (melting temperature, $T_m$ or $T_{onset}$) of the unfolding transition, with peaks corresponding to the transition points.

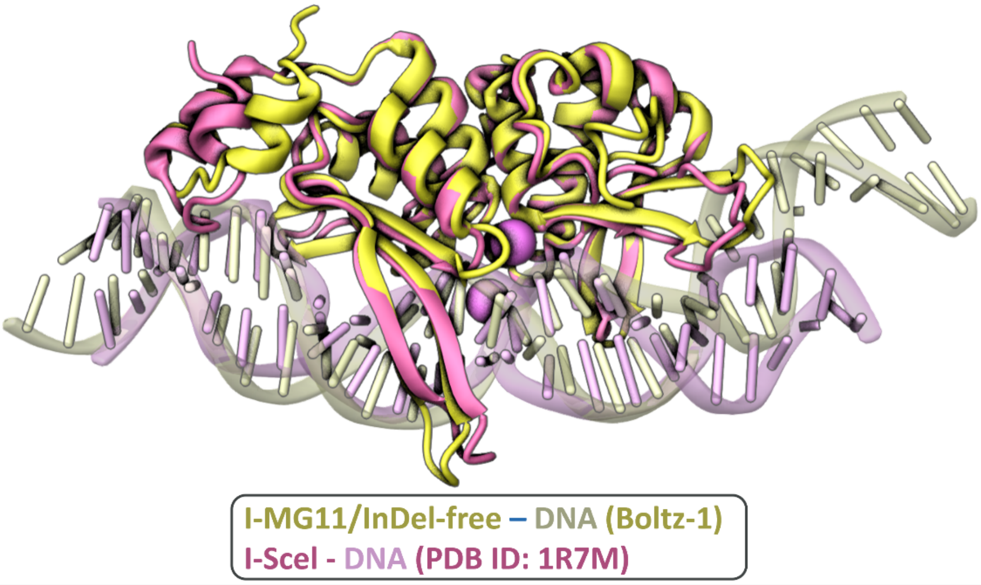

```
                            10        20        30        40        50        60        70        80
I-SceI             MKNIKKNQVMNLGPNSKLLKEYKSQLIELNIEQFEAGIGLILGDAYIRS--RDEGKTYCMQFEWKNKAYMDHVCLLYDQWVL
I-MG11             -----------MFLRGKKLQEYKSKLV-LSQRQRELLIGSLLGDGSMRIPGHNRNANFIVAHGEDQKEYVWWKYSLLKEWVL
I-MG11_No_Indels   MKNIKKNQVMNLFLRGKKLQEYKSKLV-LSQRQRELLIGSLLGDGSMRI--HNRNANFIVAHGEDQKEYVWWKYSLLKEWVL

                          90       100       110       120       130       140       150       160
I-SceI             SPPHKKERVNHLGN--LVITWGAQTFKHQAFNKLANLFIVNNKKTIPNNLVENYLTPMSLAYWFMDDGGKWDYNKNSTNKSI
I-MG11             TPPKRSIRRYHKDKTKSTISWRFLTLSHPHLTEMYQVFYRDNRKIIPNDIEKLLISPFSLAVWIMDDGSR-------NREAL
I-MG11_No_Indels   TPPKRSIRRYHKDK--STISWRFLTLSHPHLTEMYQVFYRDNRKIIPNDIEKLLISPFSLAVWIMDDGSRWDYNKNSNREAL

                         170       180       190       200       210       220       230       240
I-SceI             VLNTQSFTFEEVEYLVKGLRNKFQLNCYVKIN-----KNKPIIYIDSMSYLIFYNLIKPYLIPQMMYKLPNTISSETFLK
I-MG11             FLNTQNFSFREQERLCECLKKNFNLDATINVHSVYNGKQFYRIRINTESTRKLCCLVQEYLLPSMYYKLPFPRN-----D
I-MG11_No_Indels   FLNTQNFSFREQERLCECLKKNFNLDATINVH-----KQFYRIRINTESTRKLCCLVQEYLLPSMYYKLPFPRNSETFLD
```

**Figure S10:** Probing the effect of *in silico* removal of InDels from I-MG11 creates a mutant structurally similar I-SceI. *Top*: structural alignment of the I-MG11-InDel-free Boltz-1 model with I-SceI meganuclease (PDB ID: 1R7M) bound to their target DNA. The two enzymes and the DNA helices are shown in cartoon representations in blue and yellow colours for I-MG11 and I-SceI, respectively. *Bottom*: amino acid sequence alignment of I-MG11-InDel-free, I-MG11 and I-SceI, where α helices and β sheets secondary structures are highlighted by cylinders and arrows, respectively. InDels in I-MG11, relative to I-SceI, are reverted back in I-MG11-InDel-free, where insertions are in indicated by bold text and deletions are as hyphens (-) sign.

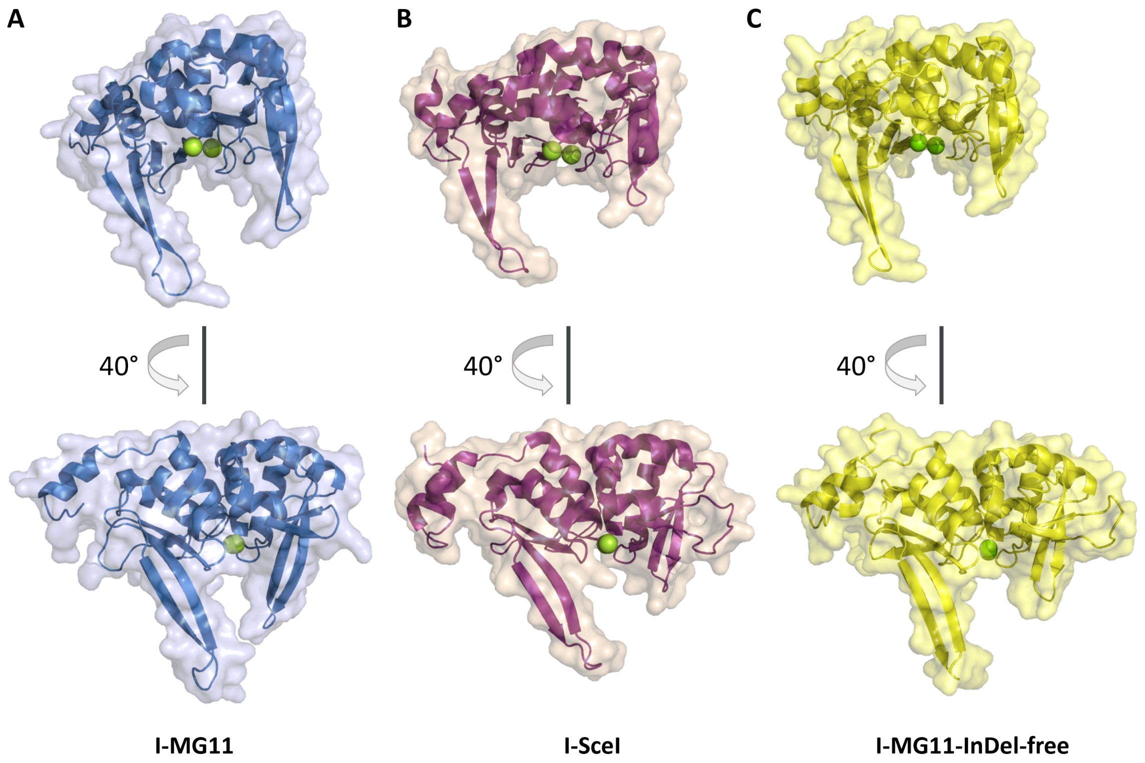

**Figure S11:** DNA-binding pocket comparison between I-MG11, I-SceI, and I-MG11-InDel-free. A more closed saddle-like DNA binding pocket in I-MG11 compared to I-SceI and I-MG11-InDel-free. **A**) Boltz-1 model of I-MG11. **B**) I-SceI structure (PDB ID: 1R7M). C) Boltz-1 model of I-MG11-InDel-free.

```
         1                                                          60
I-MG11   -------------------GTACGCGA--GCTGGGTTC----------------------
H-DreI   CAA--------------AACGTCGTAA--GTTCCGGCGC--------------------G
I-AniI   ------------------------TGA--GGAGGTTTCTCTG--------------TAAA
I-CeuI   ------ATAACGGTCCTAAGGTAGCGA--A------------------------------
I-CreI   CAA--------------AACGTCGTGA--GACAGTTTG----------------------
I-CvuI   ----TCAGAACGTC-----GTACGACG--TTCTGA-------------------------
I-DmoI   ------------------GCCTTGCCG--GGTAAGTTCC---------------GGCGCG
I-GzeII  --------------------------G--GATGGGTACCATATTGGTACAAAGGG-----
I-LtrI   --------AATGCTC----CTATACGAC-GTTTAG-------------------------
I-LtrWI  GGTTAAATAAC--------ATACTTCACTACTGG--------------------------
I-MsoI   CAG--------------AACGTCGTGA--GACAGTTCG----------------------
I-OnuI   ------------TTTCCACTTATTCAA--CCTTTTA------------------------
I-SceI   -------------------TAGGGATA--ACAGGGTAAT---------------------
I-WcaI   -------------------TAGGGATA--ACAGGGTAAT---------------------
PI-SceI  ---------------------ATCTAT--GTCGGGTGC-------GGAGAAAGAGGTAAT
```

**FigureS12:** Comparison of target sequences from different meganucleases. Alignment of the I-MG11 target site with the target sequence of previously characterized meganucleases, showing substantial differences. Mismatches are highlighted in red color. All target sequences are available in **Table S2**.

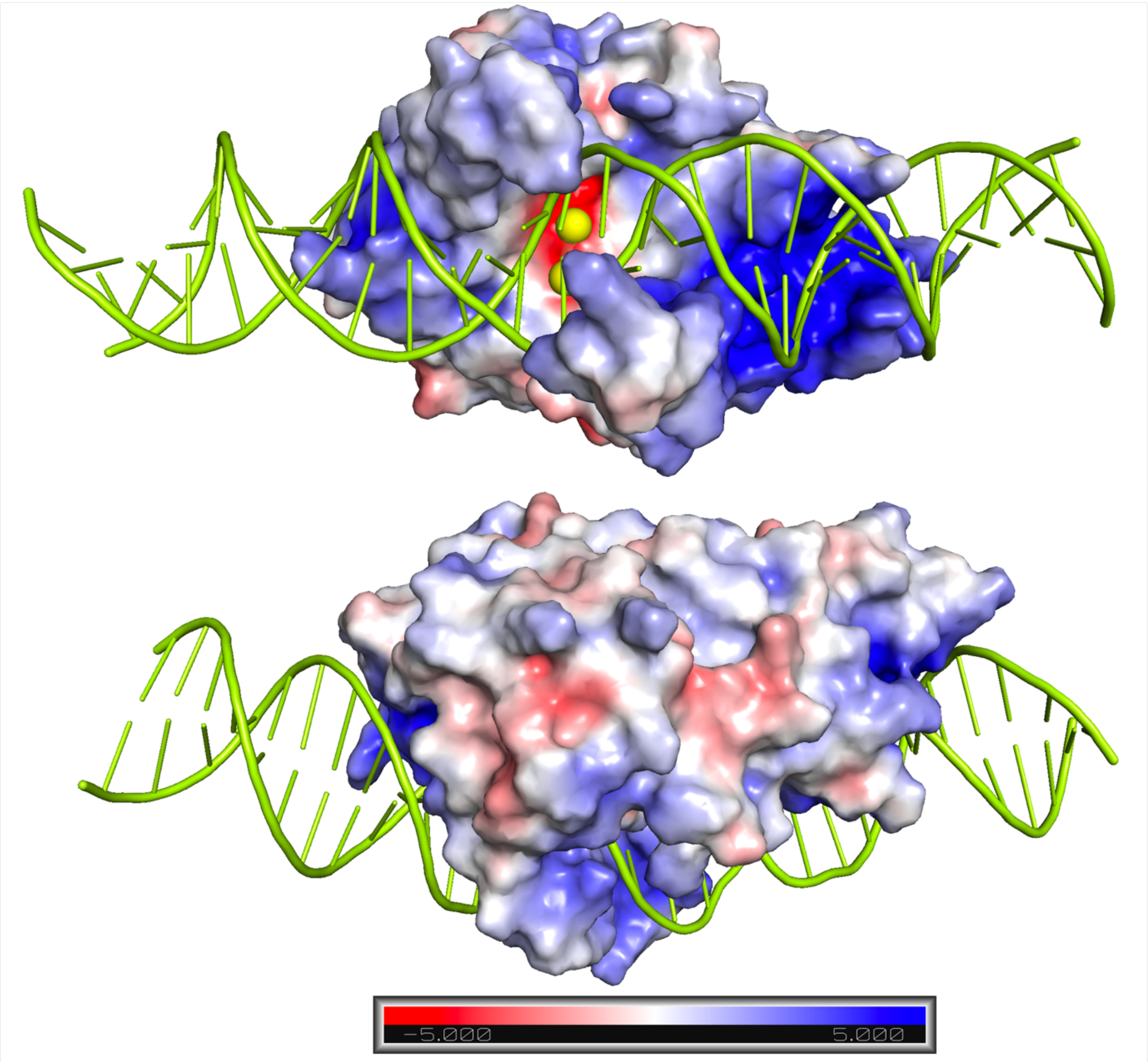

**FigureS13:** Mapping of charge distribution on the surface of I-MG11 Boltz-1 model (**Figure 7** and **Supplementary File S4**). Positively charged patches are distinguishable in the DNA-binding pocket (top) compared to the non-DNA-binding side of the protein (bottom).

#### 2. Supplementary Tables

**Table S1:** List of 16 putative LAGLIDADG single chain enzymes with average length of 215 amino acids (AA) and isoelectric point (pI) of 10.05.

| Protein | MGnify Identifier | AA | pI | AA Sequence |
|---|---|---|---|---|
| 1 | MGYP000347909486 | 217 | 10.43 | MRSRTIEEYKRGLALCAEQREALVGLLLGDAHLEGDRPGRTYRLKIEQSDRRRAYVEHLHRLFHEWVLTPPQPKI<br>VRSRGHASVNWWFQTVRHKAFQFYATQFYRDGRKCVPRLLDQMLTPRGLAYWFMDDGSRKWRQSRAVLLNT<br>QGYPKEDVEYLAQVLLQNFGLETYLRRQRDGYQIVIAGRSLERFAMLIAPYLLPEMWYKLPRVGQTQLPKR |
| 2 | MGYP000372966493 | 218 | 10.52 | MRTKEIEEQKKKLRLTEEQREVLIGILLGDAHLETQNRGRTYRLKIEQSERHKNYLFHLYEIFKEWVLTP<br>PQFRIREYPSGKKSIHWWFQTVSHGAFRFYAHQFYKEGKKRVPKLIHRWLKPRSLAYWFMDDGSIKW<br>SKSKALILNTQAFSPGDINRLIRVLNNVYQLEVYPRKQKDGVQIVISGKTFHRFVEIIKPYLLPSMIYKLPGGGQTQ<br>MPKE |
| 3 | MGYP000558734707 | 212 | 10.08 | MRSREIEEYKKGLKLTREQREVLVGLMLGDACLETQDNGRTYRLKVEQGEAQLAYIRHLYQLFAEWAQTP<br>PQAKLVRSRGRVSRNWWFQTLSHGAFRFYAHQFYRDGRKCVPRLIHRWLTPRALAYWFMDDGFVKSAQ<br>SKGVLINAQAFSLGEVQRLCQVLQERFDLKAYPRKQREGYQVYISGESYKLFRDLIEPYLIPEMRYKLPPAGQT |
| 4 | MGYP000218035350 | 209 | 10.2 | MNSKEIGQYKSKLKLSDIQRQVIVGLLLGDAHLETQNKGRTYRLKIEYSVKHSDYAMHIYELFKEWILTPPRIK<br>SDITHNNICFQTVSHAAFRFYAHQFYKDGKKSVPTKLIHKLLSPRSMAYWFMDDGSIKSRQSEGIILNTQCYG<br>EKEVEFLAKTLKRLFGLKAEKRKQKKGWQIYISGHSYETFRKIVDPYIETSMRYKIPSDRKT |
| 5 | MGYP000662234377 | 225 | 10.37 | RNYEQPIILNTKNMDKKQSKVLPESLSQIQRDVLVGLLLGDGHLETQNGGRTYRLKVEHGLKQKDYVHWLY<br>GIFSEWVPNGIYNKNRKDGSKSIGFTTRSHGTFRFYGQQFYGENQKKHIPTLIRKILTPIGMAVWFMDDGSR<br>KSLRHNTYNIHTLGYLKKDLEVMQDVLLKMFNIRTRLHKQKGKYHRIYILSESAMRFTDLIKEYVIPIQSMEHKLVT<br>KMPKE |
| 6 | MGYP000642748079 | 208 | 10.33 | MRSKEIEAYKSKLKLTKAQKEILVGTLLGDGHLETQNGRTYRLKIEHSINQKEYIDWLYSKFKQWTRTEPKMK<br>LKNGEPSTHYFNTYSHGAFRFYAQQFYVGKEKRIPKLIGKMLTPLSLAIWFMDDGSIKSSKHKSVLFHTLGYE<br>KKELEQLQFVLLEKFNLKTSLHKQKQKYWRIYIKSESMGAFKSLVGPYIITSMKYKLGNVDA |
| 7 | MGYP000621355806 | 232 | 10.22 | PRSFTRKQLIDYKNKLELTSEQKEVLIGTLLGDSSMSIRSGKPHYSVKFEQGENHKDYVDHLYQIFEPYTGSPPSM<br>RFIDSKKTRRAYWFRTYQHKHFMYYFHLFYVITPNHENVGNTSPKCKKVKIVPKNIHKYLTPRVLAYWFMDDGTF<br>H<br>CPPRSKQKSYLFSTQGFQKHEVKRLCDALKYKFNIRANVVRDKTNWRIYILKESSTTFVEPRRPHIQASSLSTNQT<br>VQAD |
| 8 | MGYP000666359522 | 219 | 9.91 | SNEQLTQYLFEITTGILLGDGNLQKPKSCRYHRLRFTQNQKRQDYVDWLFKQYKNEEFLLTSSKKLIAQNQPSE<br>FSYITSKNKKNEKGYYFQTRISSAFNKHAEIFYTENSTKKRLCSELNIFDELLTPTALAFWFMDDGTWPNKNSRN<br>FLLCTHGFTVNQVKYFSNLLNNKFGLITTVRFNKGQPVISISAKCFNKFENLILPYIQLIPSMINKFPVT |
| 9 | MGYP000097256344 | 204 | 9.85 | AASSRERDIILGTILGDAHLYSMKTDARLEIGHSEKQKEYVLYKYGELKRFVGAQPHRVKIFDPRYQKTYVQWRF<br>KTKASPLLTEFHRYFYRETKKIIPKDIASILTSPLSLAIWFMDDGGRRNDCYGLFLNTLSFTKDEHEVLRKCLKQNF<br>SLDTRLHWIQDGYRIYIPSSNAKHFCELVDPFIIPSMKYKLPYNPVTTSFAR |
| 10 | MGYP000682052720 | 204 | 9.11 | MNELQREVLIGCCLGDISIQEVSNKTHRFNMTHSIKQGEYLVHKYNIFKDLCNTPYKQRDTRYPVLYFTTLSNSD<br>IKKVTECFYNDGIRGVPQNIHEILTSPRSLAYWFMDDGTCTYNSIVKRMKVRNSTVTLCTDRYSVEEVELLISTLQ<br>NNFGINSKIKHSKSMKDRYRIYIGTKDTQKFLDIVEPFIIPSMMYKVKRPYTL |
| **11** | **MGYP00036547890<br>7** | **218** | **10.21** | **FLRGKKLQEYKSKLVLSQRQRELLIGSLLGDGSMRIPGHNRNANFIVAHGEDQKEYVWWKYSLLKEWVLTP<br>PKRSIRRYHKDKTKSTISWRFLTLSHPHLTEMYQVFYRDNRKIIPNDIEKLLISPFSLAVWIMDDGSRNREALF<br>LNTQNFSFREQERLCECLKKNFNLDATINVHSVYNGKQFYRIRINTESTRKLCCLVQEYLLPSMYYKLPFPRN** |
| 12 | MGYP000273781939 | 218 | 9.25 | RKYTKEILNRQTNSVKHHRVASIELTQEQLEVLYGSLLGDMCITKSKNLCRVCINHGGDQEAYFDHKCAIFDGLL<br>GKISKTPRYDKRTNKYYNKFAVRLLSHSTYNKLYDQLYVNGIKTLTREWLDKVTPRGLAFWFMDDGCNSGTLAT<br>NCFSLDECKTIQQWLKEVYNVDTTLQKAPNEQYIVYIKSSSRRTFYNIVIPYIIPSMLYKLNNWNLKPC |
| 13 | MGYP000292104859 | 225 | 9.55 | NPTTKYSFAKRIRSEYRIGPHNKDILSIFYGSLLGDSHAEKRKEGHGTRFSFSQESSHKSYLLWLHSIIAEKGYC<br>NPTIPVIQSRIGPGGNIRYILRFHTFTYSSLNWVHNEWYKDGSKQVPSNIEEYLTPLAIAIWIMDDGTRQGKTLKW<br>ATNAFSYKDCFLLTEVLYKKYNIKCNIHSAGKENQYVIYVMKESMPVLYHLVKDYMVSSMLYKIQELFSNENWK |
| 14 | MGYP000196790501 | 201 | 9.9 | AKERIGPHQEQVISTLVGNLLGDGHMEKRSNATRMHIHMSSRNVEYINWLHIFFSEKGYCSTEKLKLSKQIG<br>KNNKIYFSCKFRTFSFSSFNFLHDSFYVKKKKKVPEEIYNLLSPRALAIWIMDDGGKSGAGVKISTDCFCLQDVI<br>RLQTVIFEKYKITCTVQKHKSNYVIYFPKNQLSCLSKTVKPYMIPCMYYKLNNY |
| 15 | MGYP000471040085 | 193 | 10.34 | MNQTTNSQKLREYKKSLKLTQEQREALFGILLGDGHLETRDCGRTYRLLIEQTSKNERYIQHLYELFKPWVLEP<br>PKTKKRNAGVNIFFRTLSHPAFRFYGQLLYPRNAPTETTFPKIIPKNIHKYFSDRALAYWYMDDGARKGANRSG<br>KRIHTEGFEEYQVRELCQALNLYGVETTVQKQKRRPFSKKRNLES |
| 16 | MGYP000070196699 | 233 | 10.51 | MKGRSKLLSEYKNTLKLSSLQRQTLIGILLGDGCLETQNNGRTYRLLIAQSEQHALYCDHLYSIFSKWVLTPPK<br>FSKRSNGLKMKTFKTISHSSLRFYGQQFYENLAVNQNAETGFKKRRKKLPKLIHRWLSARSLAYWFMDDGSCKS<br>AT<br>SFGVILNTHNFKLCEVQTLCQILTNRWNLQCWPRKQKNKFYKKVYYQIYISGRSFKQFLKIIEPYLIDSMRYKLPPP<br>YLNKIQ |
| | **Mean** | **215** | **10.05** | |

**Table S2**:

| Enzyme | DNA Substrate (5'-->3') | **Reference** |
|---|---|---|
| I-SceI | TAGGGATAACAGGGTAAT | (1) |
| I-CreI | CAAAACGTCGTGAGACAGTTTG | (2) |
| PI-SceI | ATCTATGTCGGGTGCGGAGAAAGAGGTAAT | (3) |
| I-AniI | TGAGGAGGTTTCTCTGTAAA | (2) |
| I-CeuI | ATAACGGTCCTAAGGTAGCGAA | (4) |
| I-DmoI | GCCTTGCCGGGTAAGTTCCGGCGCG | (5) |
| H-DreI | CAAAACGTCGTAAGTTCCGGCGCG | (6) |
| I-MsoI | CAGAACGTCGTGAGACAGTTCG | (7) |
| I-WcaI | TAGGGATAACAGGGTAAT | (8) |
| I-CvuI | TCAGAACGTCGTACGACGTTCTGA | (9) |
| I-LtrWI | GGTTAAATAACATACTTCACTACTGG | PDB: 4LQ0 *(Entry by Chik et al, (2014))* |
| I-LtrI | AATGCTCCTATACGACGTTTAG | (10) |
| I-OnuI | TTTCCACTTATTCAACCTTTTA | (10) |
| I-GzeII | GGATGGGTACCATATTGGTACAAAGGG | PDB: 4EFJ *(Entry by Kulshina, N., 2012)* |
| **I-MG11** | **GTACGCGAGCTGGGTTC** | **This study** |
